# Omicron BA.2 breakthrough infection enhances cross-neutralization of BA.2.12.1 and BA.4/BA.5

**DOI:** 10.1101/2022.08.02.502461

**Authors:** Alexander Muik, Bonny Gaby Lui, Maren Bacher, Ann-Kathrin Wallisch, Aras Toker, Andrew Finlayson, Kimberly Krüger, Orkun Ozhelvaci, Katharina Grikscheit, Sebastian Hoehl, Sandra Ciesek, Özlem Türeci, Ugur Sahin

## Abstract

Recently, we reported that BNT162b2-vaccinated individuals after Omicron BA.1 breakthrough infection have strong serum neutralizing activity against Omicron BA.1, BA.2, and previous SARS-CoV-2 variants of concern (VOCs), yet less against the highly contagious Omicron sublineages BA.4 and BA.5 that have displaced previous variants. As the latter sublineages are derived from Omicron BA.2, we characterized serum neutralizing activity of COVID-19 mRNA vaccine triple-immunized individuals who experienced BA.2 breakthrough infection. We demonstrate that sera of these individuals have broadly neutralizing activity against previous VOCs as well as all tested Omicron sublineages, including BA.2 derived variants BA.2.12.1, BA.4/BA.5. Furthermore, applying antibody depletion we showed that neutralization of BA.2 and BA.4/BA.5 sublineages by BA.2 convalescent sera is driven to a significant extent by antibodies targeting the N-terminal domain (NTD) of the spike glycoprotein, whereas their neutralization by Omicron BA.1 convalescent sera depends exclusively on antibodies targeting the receptor binding domain (RBD). These findings suggest that exposure to Omicron BA.2, in contrast to BA.1 spike glycoprotein, triggers significant NTD specific recall responses in vaccinated individuals and thereby enhances the neutralization of BA.4/BA.5 sublineages. Given the current epidemiology with a predominance of BA.2 derived sublineages like BA.4/BA.5 and rapidly ongoing evolution, these findings are of high relevance for the development of Omicron adapted vaccines.

## Introduction

Emergence of the SARS-CoV-2 Omicron variant of concern (VOC) in November 2021 (*1*) can be considered a turning point in the COVID-19 pandemic. Omicron BA.1, which is significantly altered in the spike (S) glycoprotein receptor binding domain (RBD) and N-terminal domain (NTD), partially escapes previously established immunity (*2*). The loss of many epitopes (*3, 4*) drastically impaired susceptibility to neutralizing antibodies induced by wild-type strain (Wuhan-Hu-1) S glycoprotein-based vaccines or by infection with previous strains (*5–7*), necessitating a third vaccine dose to establish full immunity (*8–10*). Omicron BA.1 was displaced by the BA.2 variant, which in turn was displaced by its descendants BA.2.12.1, BA.4 and BA.5 that in the meantime dominate in many regions (*11–14*).

Antigenically, BA.2.12.1 exhibits high similarity with BA.2 but not BA.1, whereas BA.4 and BA.5 differ considerably from BA.2 and even more so from BA.1, in line with their genealogy (*15, 16*). While some amino acid changes in the RBD are shared between all Omicron sub-lineages, the alteration L452Q is only found in BA.2.12.1 and is the only residue which distinguishes its RBD from that of the BA.2 variant. The L452R and F486V alterations are BA.4/BA.5-specific, whereas S371F, T376A, D405N, and R408S are shared by BA.2 and its descendants BA.2.12.1 and BA.4/BA.5, but not BA.1 (fig. S1). These amino acid exchanges are associated with further escape from vaccine-induced neutralizing antibodies and therapeutic antibody drugs targeting the wild-type S glycoprotein (*6, 15, 17-20*). The NTDs of BA.2 and its descendants are antigenically closer to the wild-type strain and lack several amino acid changes, insertions, and deletions that occurred in BA.1 (fig S1). For instance, Δ143-145, L212I, or ins214EPE, which rendered the BA.1 variant resistant to a panel of NTD-directed monoclonal antibodies raised against the wild-type S glycoprotein, are not found in BA.2 and descendants (*21, 22*).

We and others (*10, 23*) have recently shown that Omicron BA.1 breakthrough infection of BNT162b2 vaccinated individuals augments broadly neutralizing activity against Omicron BA.1, BA.2 and previous VOCs at levels similar to those observed against SARS-CoV-2 wild-type. We showed that BA.1 breakthrough infection of triple BNT162b2-vaccinated individuals induced a robust recall response, primarily expanding memory B cells against epitopes shared broadly amongst variants, rather than inducing B cells specific to BA.1 only. Neutralization of the latest Omicron sublineages BA.4 and BA.5 was not enhanced, and geometric mean titers were rather comparable to those against the phylogenetically more distant SARS-CoV-1.

Given that Omicron BA.2 is more closely related to BA.4/BA.5 than to BA.1, we asked if BA.2 breakthrough infection would shift cross-neutralization activity more towards these most recent Omicron sublineages. We compared the neutralization of different Omicron sublineages by serum samples from three different cohorts of individuals triple-vaccinated with mRNA COVID-19 vaccines, namely from individuals with no history of SARS-CoV-2 infection and individuals that experienced breakthrough infection with either BA.1 or BA.2. In addition, we characterized the contribution of serum antibodies targeting the S glycoprotein RBD versus the NTD to Omicron sublineage neutralization. Our data will increase current understanding on Omicron immune escape mechanisms and the effects of immunization on variant cross-neutralization, and thus help guide further vaccine development.

## Results

### Cohorts and sampling

This study investigated serum samples from three cohorts: from BNT162b2 triple-vaccinated individuals who were SARS-CoV-2-naïve at the time of sampling (BNT162b2^3^, n=18), from individuals vaccinated with three doses of mRNA COVID-19 vaccine (BNT162b2/mRNA-1273 homologous or heterologous regimens) who subsequently had a breakthrough infection with Omicron at a time of BA.1 dominance (mRNA-Vax^3^ + BA.1, n=14), or from triple mRNA vaccinated individuals with a breakthrough infection at a time of BA.2 dominance (mRNA-Vax^3^ + BA.2, n=13). For convalescent cohorts, relevant intervals between key events such as the most recent vaccination and infection are provided in Fig. 1 and Table S1 to S3. Sera were derived from the biosample collections of BNT162b2 vaccine trials and from a non-interventional study researching vaccinated patients that had experienced Omicron breakthrough infection.

**Fig. 1.**
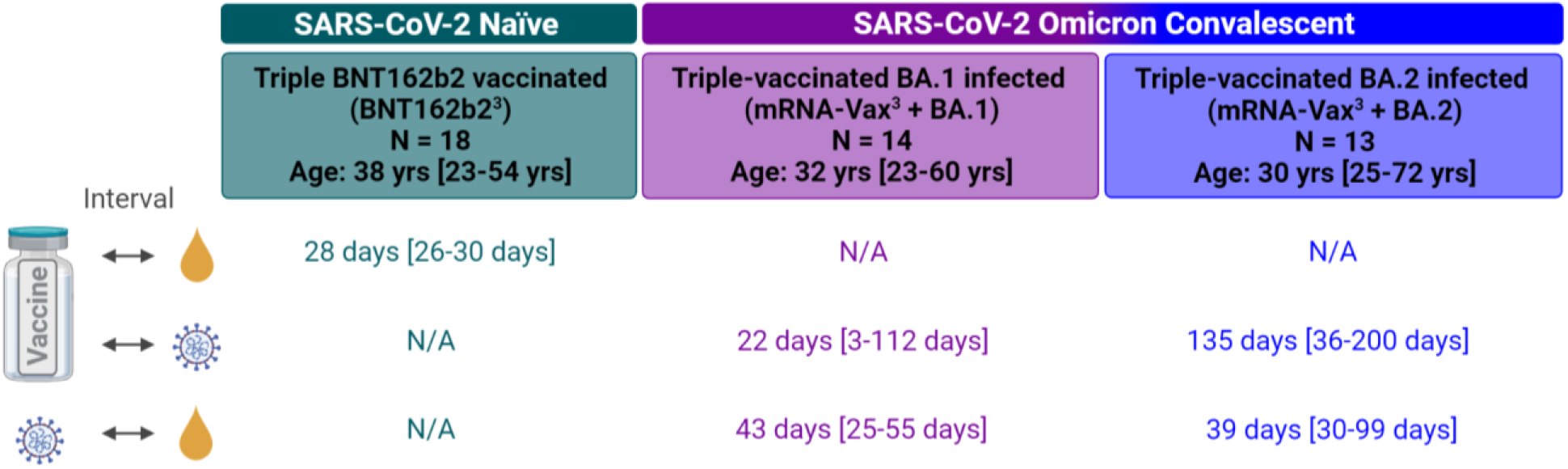
Cohorts and sampling. Serum samples (yellow droplet) were drawn from three cohorts: individuals triple-vaccinated with BNT162b2 that were SARS-CoV-2-naïve at the time of sampling (BNT162b2^3^, green), and from individuals vaccinated with three doses of mRNA COVID-19 vaccine (BNT162b2/mRNA-1273 homologous or heterologous regimens) who subsequently had a breakthrough infection with Omicron either at a time of BA.1 dominance (November 2021 to January 2022; mRNA-Vax^3^ + BA.1, purple) or at a time of BA.2 dominance (March to May 2022; mRNA-Vax^3^ + BA.2, blue). For convalescent cohorts, relevant intervals between key events such as the most recent vaccination, SARS-CoV-2 infection, and serum isolation are indicated. All values specified as median-range. The age/gender composition of cohorts is further detailed in Tables S1 to S3. Data for the reference cohorts BNT162b2^3^ and mRNA-Vax^3^ + BA.1 were previously published (*10*), except for newly generated BA.2.12.1 neutralization data. N/A, not applicable; Schematic was created with BioRender.com

A subset of the samples included in this study had also been used in our previous investigation of the effect of BA.1 breakthrough infection on serum neutralizing activity and memory B cell repertoire (Quandt et al. (*10*)).

### Omicron BA.2 breakthrough infection of triple mRNA-vaccinated individuals induces broad neutralization of VOCs including Omicron BA.4/BA.5

Neutralizing activity of immune sera was tested in a well-characterized pseudovirus neutralization test (pVNT) (*24, 25*) by determining 50% pseudovirus neutralization (pVN_50_) geometric mean titers (GMTs) with pseudoviruses bearing the S glycoproteins of the SARS-CoV-2 wild-type strain, or Alpha, Beta, Delta, Omicron BA.1, BA.2, and the BA.2-derived sublineages BA.2.12.1, BA.4 and BA.5. As BA.4 and BA.5 share an identical S glycoprotein sequence, we refer to them as BA.4/5 in the context of the pVNT. In addition, we assayed SARS-CoV (herein referred to as SARS-CoV-1) to detect potential pan-Sarbecovirus neutralizing activity (*26*). As an orthogonal test system, we used a live SARS-CoV-2 neutralization test (VNT) that analyzes neutralization during multicycle replication of authentic virus (SARS-CoV-2 wild-type strain and VOCs including BA.4, except Omicron BA.2.12.1) with the antibodies present during the entire test period.

In the pVNT, sera from all three cohorts robustly neutralized the wild-type strain, Alpha, Beta, Delta VOCs as well as Omicron BA.1 and BA.2 lineages with neutralization activity being more pronounced in the breakthrough infected individuals, particularly in the BA.1 breakthrough infection cohort (mRNA-Vax^3^ + BA.1). However, serum neutralizing activity of BNT162b2 triple-vaccinated SARS-CoV-2 naïve (BNT162b2^3^) and mRNA-Vax^3^ + BA.1 individuals against BA.2.12.1 was significantly reduced compared to wild-type (p<0.05) and even more so for BA.4/5 (p<0.001; >5-fold compared to the wild-type strain) (Fig. 2a, Table S4 and S5). In contrast, sera from triple mRNA-vaccinated individuals with Omicron BA.2 breakthrough infection (mRNA-Vax^3^ + BA.2) neutralized the BA.2.12.1 pseudovirus as robustly as the wild-type strain. Neutralization of BA.4/5 was broadly similar to that of BA.2.12.1, and the reduction relative to the wild-type strain significant (p<0.05) yet less pronounced (∼2.5-fold) as compared to the two other cohorts.

**Fig. 2.**
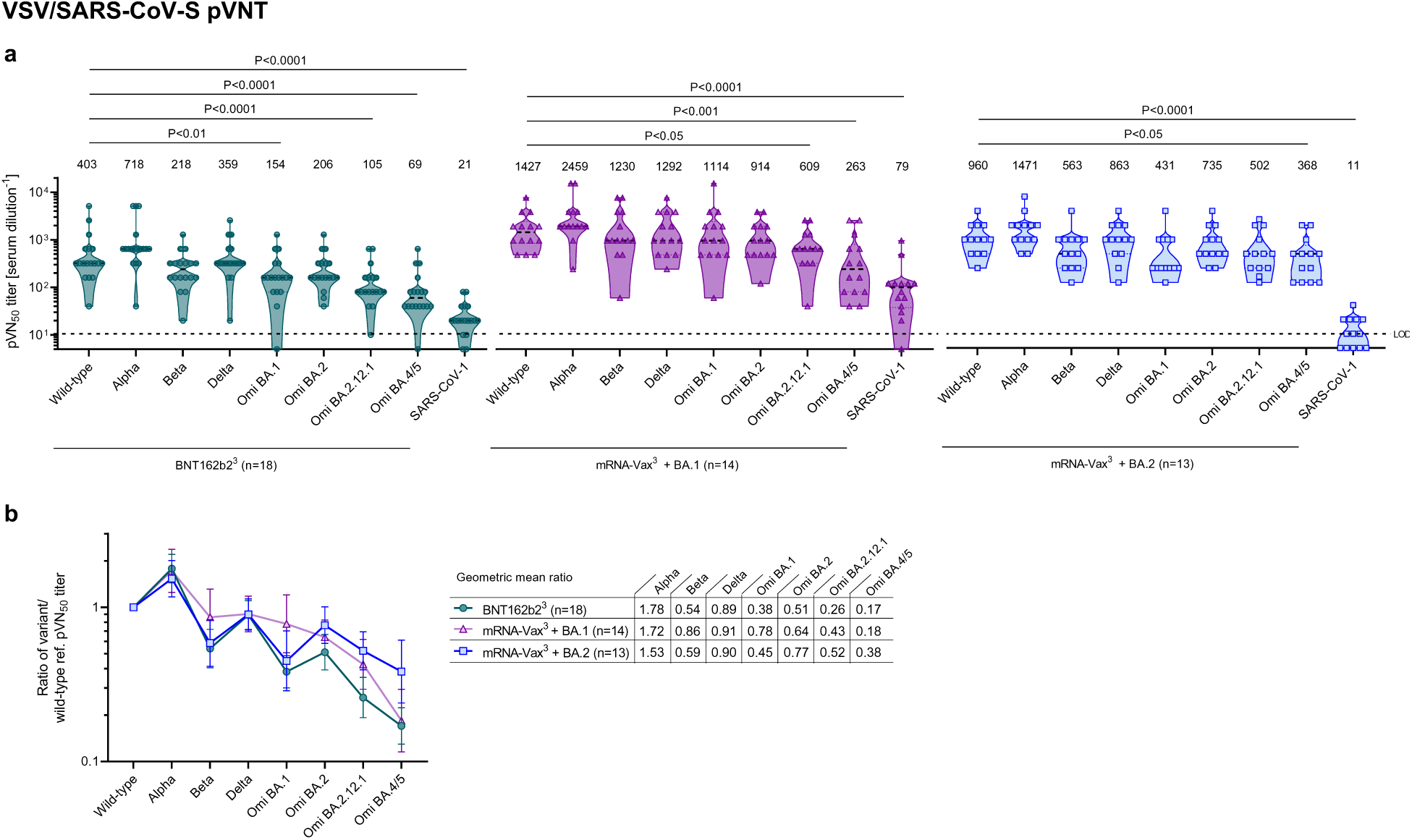
Omicron BA.2 breakthrough infection of triple mRNA vaccinated individuals induces broad neutralization of SARS-CoV-2 variant pseudoviruses including Omicron BA.4/5. Cohorts and serum sampling as described in Fig. 1. (a) 50% pseudovirus neutralization (pVN_50_) geometric mean titers (GMTs) against the indicated SARS-CoV-2 variants of concern (VOCs) or SARS-CoV-1 pseudoviruses. Data for the reference cohorts BNT162b2^3^ and mRNA-Vax^3^ + BA.1 were previously published (*10*), except for newly generated BA.2.12.1 neutralization data. Values above violin plots represent group GMTs. (b) SARS-CoV-2 VOC pVN_50_ GMTs normalized against the wild-type strain pVN_50_ GMT (ratio VOC to wild-type). Group geometric mean ratios with 95% confidence intervals are shown. Serum was tested in duplicate. For titer values below the limit of detection (LOD), LOD/2 values were plotted. The non-parametric Friedman test with Dunn’s multiple comparisons correction was used to compare the wild-type strain neutralizing group GMTs with titers against the indicated variants and SARS-CoV-1. Multiplicity-adjusted p values are shown.

To compare the cohorts with regard to neutralization breadth irrespective of the magnitude of antibody titers, we normalized the VOC pVN_50_ GMTs against the wild-type strain. The ratios showed that BA.4/5 cross-neutralization was substantially stronger in mRNA-Vax^3^ + BA.2 (GMT ratio 0.38) as compared to mRNA-Vax^3^ + BA.1 and BNT162b2^3^ sera (GMT ratios 0.18 and 0.17) (Fig. 2b). Similarly, cross-neutralization of Omicron BA.2.12.1 by mRNA-Vax^3^ + BA.2 sera (GMT ratio 0.52) was stronger than by mRNA-Vax^3^ + BA.1 sera (GMT ratio 0.43), and even more so than by BNT162b2^3^ sera (GMT ratio 0.26).

A separate analysis including only the BNT162b2 vaccinated individuals within those three cohorts confirmed that BA.2 breakthrough infection is associated with considerable BA.4/5 cross-neutralization (BA.4/5 to wild-type GMT ratio 0.42), whereas after BA.1 breakthrough infection pVN_50_ GMTs against BA.4/5 were ∼6-fold lower than those against wild-type (i.e., GMT ratio 0.17) (fig. S2a-c). Cross-neutralization of BA.2 and BA.2.12.1 by sera of the BA.1 or BA.2 convalescents was superior to that of BNT162b2 triple-vaccinated SARS-CoV-2 naïve individuals.

The authentic live SARS-CoV-2 virus neutralization assay provided VOC neutralizing titers that strongly correlated with those from the pVNT assay (fig. S3) and largely confirmed the major findings in Fig. 2. Again in this assay, 50% virus neutralization (VN_50_) GMT against Omicron BA.2 in BNT162b2^3^ sera was strongly reduced compared to that against wild-type (p<0.0001), whereas sera from both convalescent groups exhibited strong neutralizing activity, with VN_50_ GMTs comparable to those against the wild-type strain (Fig. 3a). Reduction of neutralizing activity against Omicron BA.4 was less pronounced in the BA.2 convalescent cohort as compared to BNT162b2^3^ and mRNA-Vax^3^ + BA.1 cohorts (VN_50_ GMTs ∼2.5-fold as compared to ∼15-fold and 5-fold lower than against the wild-type strain, respectively).

**Fig. 3.**
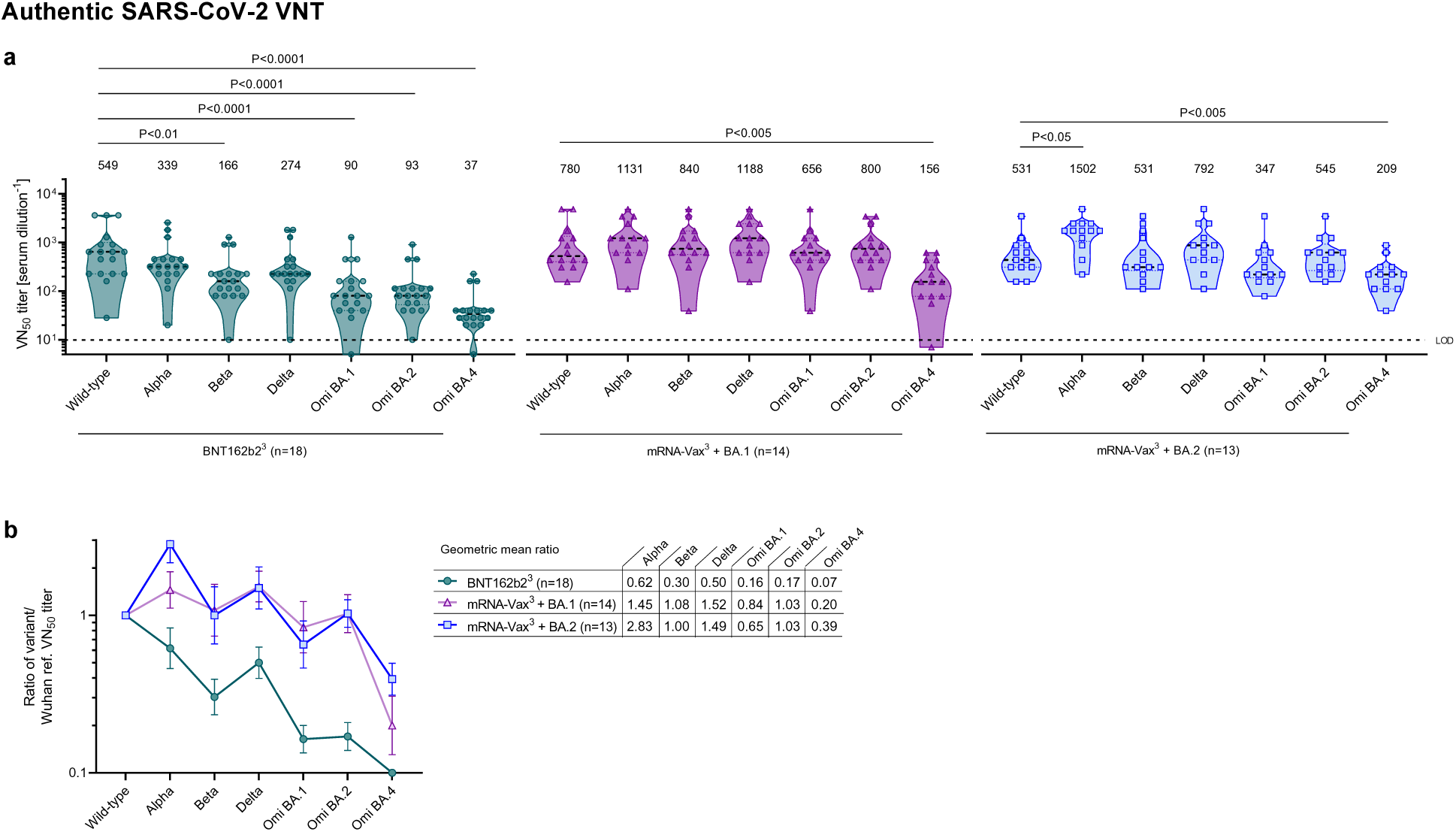
Omicron BA.2 breakthrough infection of previously vaccinated individuals induces broad neutralization of authentic live SARS-CoV-2 variants including Omicron BA.4/5. Cohorts and serum sampling as described in Fig. 1. (a) 50% virus neutralization (VN_50_) geometric mean titers (GMTs) against the indicated SARS-CoV-2 variants of concern (VOCs). Data for the reference cohorts BNT162b2^3^ and mRNA-Vax^3^ + BA.1 were previously published (*10*). Values above violin plots represent group GMTs. (b) SARS-CoV-2 VOC VN_50_ GMTs normalized against the wild-type strain VN_50_ GMT (ratio VOC to wild-type). Group geometric mean ratios with 95% confidence intervals are shown. Serum was tested in duplicate. For titer values below the limit of detection (LOD), LOD/2 values were plotted. The non-parametric Friedman test with Dunn’s multiple comparisons correction was used to compare the wild-type strain neutralizing group GMTs with titers against the indicated variants and SARS-CoV-1. Multiplicity-adjusted p values are shown.

In line with the pVNT data, magnitude-independent analyses via the calculated ratios of VOC VN_50_ GMTs against the wild-type strain showed that BA.4 cross-neutralization was stronger in the mRNA-Vax^3^ + BA.2 cohort (GMT ratio 0.39) as compared to the mRNA-Vax^3^ + BA.1 (GMT ratio 0.20) and BNT162b2^3^ (GMT ratio 0.07) cohorts (Fig. 3b) and similarly so within the sub-cohort of BNT162b2 triple-vaccinated individuals (fig. S2d-f).

In aggregate, these data demonstrate that Omicron BA.2 breakthrough infection of vaccinated individuals is associated with broad neutralizing activity against all tested Omicron-sublineages and previous SARS-CoV-2 VOCs. In particular, our data indicate that breakthrough infection with BA.2 is more effective (∼2-fold higher cross neutralization) than that with BA.1 at refocusing neutralizing antibody responses towards the BA.4/BA.5 S glycoprotein.

Neutralization of Omicron BA.2 and BA.4/5 by sera of triple mRNA vaccinated BA.2 convalescent individuals is mediated to a large extent by NTD-targeting antibodies

To dissect the role of serum antibodies binding either to the RBD or the NTD of the S glycoprotein for neutralization of SARS-CoV-2 wild-type, Omicron BA.1, BA.2, and BA.4/5, we depleted those antibody fractions separately from sera of the three cohorts (n=6 each, fig. S4a, Table S10). We used the SARS-CoV-2 wild-type strain S glycoprotein RBD and NTD baits for depletion, as VOC breakthrough infections have been demonstrated to predominantly elicit recall responses recognizing epitopes conserved across known VOCs (*10, 23, 27*).

The depletion experiments removed >97% of all RBD-binding antibodies and >74% of all NTD-binding antibodies (fig. S4b). Depleted sera were subsequently tested in pVNT assays. RBD-antibody depletion strongly diminished neutralizing activity against the wild-type strain in sera from all cohorts, whereas neutralizing activity was mostly retained (>80% remaining activity) upon depletion of NTD-binding antibodies (Fig. 4a, Table S11). Neutralization of Omicron BA.1 was completely abrogated upon depletion of RBD-binding antibodies and largely unaffected by NTD-binding antibody depletion. For neutralization of BA.2, RBD-antibody depletion almost completely abolished neutralizing activity of mRNA-Vax^3^ + BA.1 sera (about 2% residual neutralization activity). The reduction of neutralizing titers for BNT162b2^3^ and particularly mRNA-Vax^3^ + BA.2 sera was less severe with ∼12 and ∼24% remaining neutralizing activity, respectively. In contrast, depletion of NTD-binding antibodies did not considerably impact the neutralizing activity of BNT162b2^3^ and mRNA-Vax^3^ + BA.1 sera (∼91 and ∼99% of undepleted control, respectively), while neutralizing activity of mRNA-Vax^3^ + BA.2 sera was reduced to ∼50%. A similar pattern was seen following RBD-antibody depletion for neutralization of BA.4/5, with strongly reduced neutralizing activity of mRNA-Vax^3^ + BA.1 sera (∼3% residual activity) versus less severe reductions for BNT162b2^3^ and mRNA-Vax^3^ + BA.2 sera (∼20 and ∼26% remaining activity, respectively). Depletion of NTD-binding antibodies had a larger impact for BA.4/5 neutralization compared to BA.2, with remaining neutralizing activity of BNT162b2^3^ and mRNA-Vax^3^ + BA.1 sera of ∼70 and ∼90% respectively, again with the strongest effect (∼48% of undepleted control) of mRNA-Vax^3^ + BA.2 sera.

**Fig. 4.**
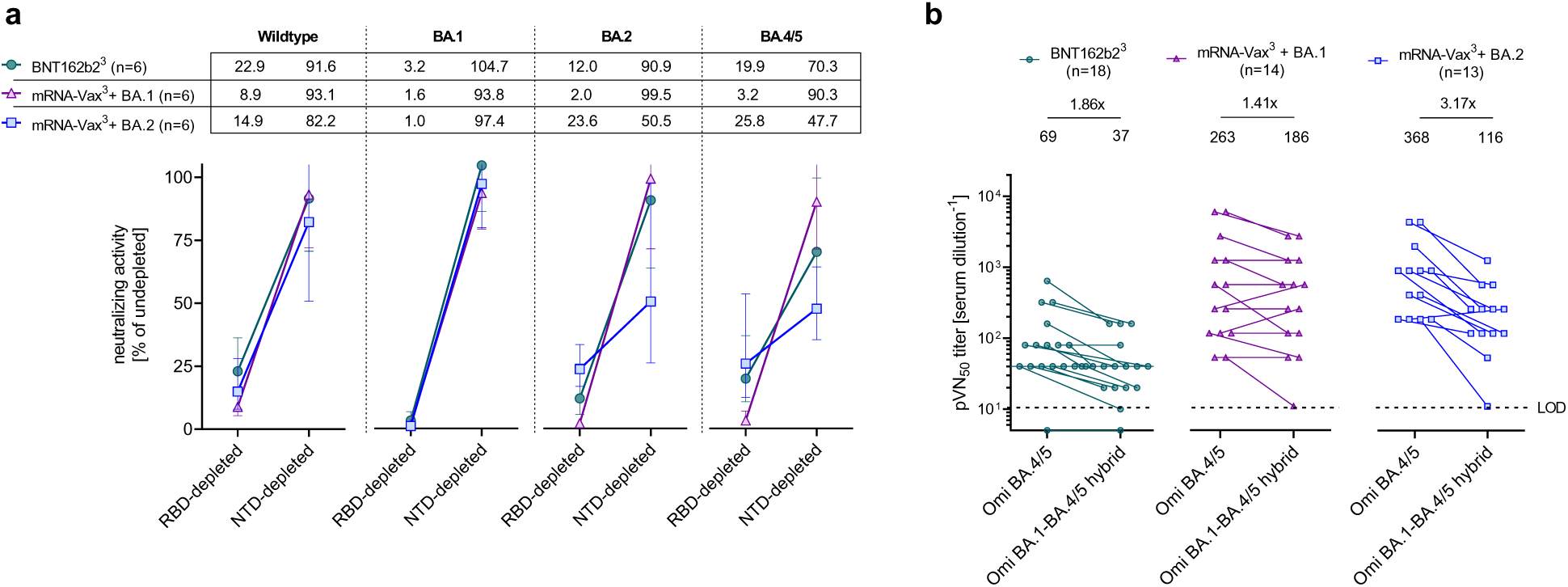
Neutralization of Omicron BA.2 and BA.4/5 by sera of triple mRNA vaccinated BA.2 convalescent individuals is mediated to a large extent by NTD-targeting antibodies. Cohorts and serum sampling as described in Fig. 1. (a) Serum samples (n=6 per cohort) were depleted of RBD- or NTD-binding antibodies. Relative neutralizing activity of RBD- and NTD-depleted sera (pVN_50_ titers of undepleted control sera were set to 100%) against the wild-type strain, BA.1, BA.2, and BA.4/5 was calculated and group geometric mean with 95% confidence intervals are shown. (b) 50% pseudovirus neutralization (pVN_50_) geometric mean titers (GMTs) against Omicron BA.4/5 and Omicron BA.1-BA.4/5 hybrid pseudoviruses. Numbers above plots indicate group geometric mean titers (GMTs) and fold-change in GMTs between BA.4/5 and the hybrid pseudovirus. For titer values below the limit of detection (LOD), LOD/2 values are plotted.

As an orthogonal approach we assessed the neutralizing activity of sera from those 3 cohorts of vaccinated individuals against a pseudovirus harboring an engineered hybrid S glycoprotein consisting of the Omicron BA.1 N-terminus including the NTD (amino acids 1-338) and the BA.4/5 C terminus including the RBD.

The pVN_50_ GMT against the Omicron BA.1-BA.4/5 hybrid pseudovirus in sera from BNT162b2^3^ was moderately below (1.86-fold) the GMT for the BA.4/5 pseudovirus, and in the BA.1 convalescents the GMT was only marginally affected (<1.5-fold reduction) (Fig. 4b, Tables S4 and S5). In contrast, in BA.2 convalescent sera titers against the hybrid pseudovirus were considerably lower than those against the BA.4/5 pseudovirus (>3-fold reduction of GMT) (Fig. 4b and Table S6), suggesting that substantial neutralizing activity can be attributed to NTD epitopes that are shared between Omicron BA.2 and BA.4/5.

In aggregate the data obtained in both experiments indicate that across all these VOCs RBD-binding antibodies have a major contribution to neutralization. Another key finding is that exposure to BA.1 (that differs substantially from previous VOCs in its NTD; fig. S1) boosts recall responses of vaccine-induced neutralizing antibodies that primarily bind the RBD, whereas exposure to BA.2 S glycoprotein (with an NTD closer related to previous VOCs) can build on existing memory and elicits a considerable recall of NTD-targeting antibodies that in turn contributes substantially to the neutralization of BA.2 and BA.4/5.

## Discussion

Recent studies have demonstrated that Omicron BA.1 breakthrough infection in individuals vaccinated with mRNA vaccines BNT162b2 or mRNA-1273 or an inactivated virus vaccine boosts serum neutralizing titers against VOCs including BA.2 (*10, 15, 23*), but not against BA.2.12.1 or BA.4/BA.5. The immune escape has been attributed to boosting of pre-existing neutralizing antibody responses that recognize epitopes shared between the SARS-CoV-2 wild-type strain and Omicron BA.1 but are in part absent in BA.2.12.1, BA.4, and BA.5 due to alterations at key residues including L452Q/L452R, and F486V (*15*).

In our current study, we report that BA.2 breakthrough infection is associated with broadly neutralizing activity including BA.2 and its descendants BA.2.12.1, BA.4 and BA.5. These findings are in agreement with recent publications (*19, 28*) and suggest that the higher sequence similarity of BA.2 with BA.2.12.1 and BA.4/5 in the S glycoprotein RBD as well as the NTD drives more efficient cross-neutralization as compared to breakthrough infections with the antigenically more distant BA.1 variant. In particular, BA.1 breakthrough infection may not elicit a strong recall of NTD-specific memory B cells owing to the substantial alterations within the BA.1 NTD (fig. S1) given that breakthrough infection with heterologous SARS-CoV-2 strains primarily expands a memory B cell repertoire against conserved S glycoprotein epitopes (*10, 23*). Our data obtained in antibody-depletion and hybrid pseudovirus experiments show that NTD-binding antibodies have a substantial contribution to neutralizing activity against Omicron BA.4/5 in triple-vaccinated BA.2 convalescent sera, whereas neutralizing activity in BA.1 convalescent sera largely relies on RBD-binding antibodies. This finding is consistent with the observation that NTD-binding antibodies isolated from BA.2 breakthrough infected individuals do not neutralize BA.1 (*29*). Together these important findings extend our knowledge on how vaccinations and boosters with the current wild-type strain-based vaccines together with breakthrough infections with the various VOCs shape the immunity patterns within the population and are material to inform further vaccine development and adaptation in response to current and emerging VOCs.

Our findings are based on retrospective analyses of samples derived from different studies, using relatively small samples sizes and cohorts that are not fully aligned in terms of intervals between vaccine doses, intervals between the most recent vaccine dose and infection, and demographic characteristics such as age and sex of individuals. While the SARS-CoV-2 naïve cohort was triple-vaccinated with BNT162b2, and the Omicron breakthrough cohorts were triple-vaccinated with BNT162b2 or mRNA-1273, or a heterologous regimen of the mRNA COVID-19 vaccines, key findings held true when only looking at a BNT162b2-vaccinated subset. Studies investigating long-lived plasma cell, memory B cell, and T cell immunity in cohorts with additional subjects could provide further insights into the mechanisms underlying the broad neutralizing activity associated with Omicron BA.2 breakthrough infection and corroborate our findings.

Notwithstanding the importance of vaccination with currently approved wild-type-strain based vaccines such as BNT162b2 that offer effective protection from severe disease by current VOCs including Omicron BA.1 and BA.2 (*30, 31*), our findings highlight that consideration of rapidly evolving epidemiological landscapes and newly emerging SARS-CoV-2 variants is of high importance for guiding vaccine adaptation programs. For instance, while the efficacy of vaccine adaptation to the BA.1 strain S glycoprotein sequence is currently under investigation in clinical trials, our data suggest that further benefit may be derived from a vaccine adapted to the sequence of BA.2 or descendants.

## Materials and Methods

### Study design, recruitment of participants and sample collection

The objective of this study was to investigate the effect of Omicron BA.2 breakthrough infection on the cross-variant neutralization capacity of human sera. We compared immune responses in triple-mRNA (BNT162b2/mRNA-1273)-vaccinated individuals with a confirmed subsequent SARS-CoV-2 breakthrough infection in a period of Omicron BA.2 lineage-dominance in Germany (March to May 2022; mRNA-Vax^3^ + BA.2), to that of triple-mRNA-vaccinated individuals with a confirmed subsequent SARS-CoV-2 breakthrough infection in a period of Omicron BA.1 lineage-dominance (November 2021 to mid-January 2022; mRNA-Vax^3^ + BA.1) (*1, 2*) and triple-BNT162b2-vaccinated individuals that were SARS-CoV-2-naïve (nucleocapsid seronegative) at the time of sample collection (BNT162b2^3^). Serum neutralizing capability was characterized using pseudovirus and live SARS-CoV-2 neutralization assays. Data for the reference cohorts BNT162b2^3^ and mRNA-Vax^3^ + BA.1 were previously published (*10*), except for newly generated BA.2.12.1 neutralization data. Cross-neutralization of variants was further characterized in smaller sub-cohorts after depletion of either wild-type S glycoprotein NTD- or RBD-targeted neutralizing antibodies.

Individuals from the BNT162b2^3^ cohort provided informed consent as part of their participation in the Phase 2 trial BNT162-17 (NCT05004181). Participants from the mRNA-Vax^3^ + Omi BA.1 and mRNA-Vax^3^ + BA.2 cohorts were recruited from University Hospital, Goethe University Frankfurt as part of a non-interventional study (protocol approved by the Ethics Board of the University Hospital [No. 2021-560]) researching patients that had experienced Omicron breakthrough infection following vaccination for COVID-19. Omicron BA.1 infections were confirmed with variant-specific PCR. The infections of 4 BA.1 convalescent participants in this study were further characterized by genome sequencing. In all 4 cases, genome sequencing confirmed Omicron BA.1 infection (Table S3).

Demographic and clinical data for all participants and sampling timepoints are provided (Tables S1 to S3, and Fig. 1). All participants had no documented history of SARS-CoV-2 infection prior to vaccination. Participants were free of symptoms at the time of blood collection. Serum was isolated by centrifugation of drawn blood at 2000 x g for 10 minutes and cryopreserved until use.

### VSV-SARS-CoV-2 S variant pseudovirus generation

A recombinant replication-deficient vesicular stomatitis virus (VSV) vector that encodes green fluorescent protein (GFP) and luciferase instead of the VSV-glycoprotein (VSV-G) was pseudotyped with SARS-CoV-1 S glycoprotein (UniProt Ref: P59594) and with SARS-CoV-2 S glycoprotein derived from either the Wuhan-Hu-1 reference strain (NCBI Ref: 43740568), the Alpha variant (alterations: Δ69/70, Δ144, N501Y, A570D, D614G, P681H, T716I, S982A, D1118H), the Beta variant (alterations: L18F, D80A, D215G, Δ242–244, R246I, K417N, E484K, N501Y, D614G, A701V), the Delta variant (alterations: T19R, G142D, E156G, Δ157/158, K417N, L452R, T478K, D614G, P681R, D950N), the Omicron BA.1 variant (alterations: A67V, Δ69/70, T95I, G142D, Δ143-145, Δ211, L212I, ins214EPE, G339D, S371L, S373P, S375F, K417N, N440K, G446S, S477N, T478K, E484A, Q493R, G496S, Q498R, N501Y, Y505H, T547K, D614G, H655Y, N679K, P681H, N764K, D796Y, N856K, Q954H, N969K, L981F),the Omicron BA.2 variant (alterations: T19I, Δ24-26, A27S, G142D, V213G, G339D, S371F, S373P, S375F, T376A, D405N, R408S, K417N, N440K, S477N, T478K, E484A, Q493R, Q498R, N501Y, Y505H, D614G, H655Y, N679K, P681H, N764K, D796Y, Q954H, N969K), the Omicron BA.2.12.1 variant (alterations: T19I, Δ24-26, A27S, G142D, V213G, G339D, S371F, S373P, S375F, T376A, D405N, R408S, K417N, N440K, L452Q, S477N, T478K, E484A, Q493R, Q498R, N501Y, Y505H, D614G, H655Y, N679K, P681H, S704L, N764K, D796Y, Q954H, N969K), the Omicron BA.4/5 variant (alterations: T19I, Δ24-26, A27S, Δ69/70, G142D, V213G, G339D, S371F, S373P, S375F, T376A, D405N, R408S, K417N, N440K, L452R, S477N, T478K, E484A, F486V, Q498R, N501Y, Y505H, D614G, H655Y, N679K, P681H, N764K, D796Y, Q954H, N969K), or an artificial Omicron BA.1-BA.4/5 hybrid S glycoprotein (alterations: A67V, Δ69/70, T95I, G142D, Δ143-145, Δ211, L212I, ins214EPE, G339D, S371F, S373P, S375F, T376A, D405N, R408S, K417N, N440K, L452R, S477N, T478K, E484A, F486V, Q498R, N501Y, Y505H, D614G, H655Y, N679K, P681H, N764K, D796Y, Q954H, N969K) according to published pseudotyping protocols (*3*). A diagram of SARS-CoV-2 S glycoprotein alterations is shown in fig. S5a and a separate alignment of S glycoprotein alterations in Omicron sub-lineages is displayed in fig. S1.

In brief, HEK293T/17 monolayers (ATCC® CRL-11268™) cultured in Dulbecco’s modified Eagle’s medium (DMEM) with GlutaMAX™ (Gibco) supplemented with 10% heat-inactivated fetal bovine serum (FBS [Sigma-Aldrich]) (referred to as medium) were transfected with Sanger sequencing-verified SARS-CoV-1 or variant-specific SARS-CoV-2 S expression plasmid with Lipofectamine LTX (Life Technologies) following the manufacturer’s instructions. At 24 hours after transfection, the cells were infected at a multiplicity of infection (MOI) of three with VSV-G complemented VSVΔG vector. After incubation for 2 hours at 37 °C with 7.5% CO_2_, cells were washed twice with phosphate buffered saline (PBS) before medium supplemented with anti-VSV-G antibody (clone 8G5F11, Kerafast Inc.) was added to neutralize residual VSV-G-complemented input virus. VSV-SARS-CoV-2-S pseudotype-containing medium was harvested 20 hours after inoculation, passed through a 0.2 µm filter (Nalgene) and stored at -80 °C. The pseudovirus batches were titrated on Vero 76 cells (ATCC® CRL-1587™) cultured in medium. The relative luciferase units induced by a defined volume of a SARS-CoV-2 wild-type strain S glycoprotein pseudovirus reference batch previously described in Muik et al., 2021 (*4*), that corresponds to an infectious titer of 200 transducing units (TU) per mL, was used as a comparator. Input volumes for the SARS-CoV-2 variant pseudovirus batches were calculated to normalize the infectious titer based on the relative luciferase units relative to the reference.

### Pseudovirus neutralization assay

Vero 76 cells were seeded in 96-well white, flat-bottom plates (Thermo Scientific) at 40,000 cells/well in medium 4 hours prior to the assay and cultured at 37 °C with 7.5% CO_2_. Each individual serum was serially diluted 2-fold in medium with the first dilution being 1:5 (SARS-CoV-2-naïve triple BNT162b2 vaccinated; dilution range of 1:5 to 1:5,120) or 1:30 (triple vaccinated after subsequent Omicron BA.1 or BA.2 breakthrough infection; dilution range of 1:30 to 1:30,720). In the case of the SARS-CoV-1 pseudovirus assay, the serum of all individuals was initially diluted 1:5 (dilution range of 1:5 to 1:5,120). VSV-SARS-CoV-2-S/VSV-SARS-CoV-1-S particles were diluted in medium to obtain 200 TU in the assay. Serum dilutions were mixed 1:1 with pseudovirus (n=2 technical replicates per serum per pseudovirus) for 30 minutes at room temperature before being added to Vero 76 cell monolayers and incubated at 37 °C with 7.5% CO_2_ for 24 hours. Supernatants were removed and the cells were lysed with luciferase reagent (Promega). Luminescence was recorded on a CLARIOstar® Plus microplate reader (BMG Labtech), and neutralization titers were calculated as the reciprocal of the highest serum dilution that still resulted in 50% reduction in luminescence. For depletion studies resolution with regards to neutralization titers was increased, in order to discriminate smaller than 2-fold differences on an individual serum level. Neutralization titers were determined by generating a 4-parameter logistical (4PL) fit of the percent neutralization at each serial serum dilution. The 50% pseudovirus neutralization (pVN_50_) titer was reported as the interpolated reciprocal of the dilution yielding a 50% reduction in luminescence. Results for all pseudovirus neutralization experiments were expressed as geometric mean titers (GMT) of duplicates. If no neutralization was observed, an arbitrary titer value of half of the limit of detection [LOD] was reported. Tables of the neutralization titers are provided (Tables S4 to S6, Table S11). SARS-CoV-2 wild-type strain, and Alpha, Beta, Delta, BA.1, BA.4/5 VOC, as well as SARS-CoV-1 pseudovirus neutralizing GMTs for the SARS-CoV-2 naïve BNT162b2 triple-vaccinated cohort and the triple-vaccinated BA.1 convalescent cohort were previously reported in Quandt. et al. (*10*). Only the BA.2.12.1 neutralization data was newly generated from serum samples for this study.

### Live SARS-CoV-2 neutralization assay

SARS-CoV-2 virus neutralization titers were determined by a microneutralization assay based on cytopathic effect (CPE) at VisMederi S.r.l., Siena, Italy. In brief, heat-inactivated serum samples from individuals were serially diluted 1:2 (starting at 1:10; n=2 technical replicates per serum per virus) and incubated for 1 hour at 37 °C with 100 TCID_50_ of the live wild-type-like SARS-CoV-2 virus strain 2019-nCOV/ITALY-INMI1 (GenBank: MT066156), Alpha virus strain nCoV19 isolate/England/MIG457/2020 (alterations: Δ69/70, Δ144, N501Y, A570D, D614G, P681H, T716I, S982A, D1118H), Beta virus strain nCoV19 isolate/England ex-SA/HCM002/2021 (alterations: D80A, D215G, Δ242–244, K417N, E484K, N501Y, D614G, A701V), sequence-verified Delta strain isolated from a nasopharyngeal swab (alterations: T19R, G142D, E156G, Δ157/158, L452R, T478K, D614G, P681R, R682Q, D950N), Omicron BA.1 strain hCoV-19/Belgium/rega-20174/2021 (alterations: A67V, Δ69/70, T95I, G142D, Δ143-145, Δ211, L212I, ins214EPE, G339D, S371L, S373P, S375F, K417N, N440K, G446S, S477N, T478K, E484A, Q493R, G496S, Q498R, N501Y, Y505H, T547K, D614G, H655Y, N679K, P681H, N764K, D796Y, N856K, Q954H, N969K, L981F), sequence-verified Omicron BA.2 strain (alterations:T19I, Δ24-26, A27S, V213G, G339D, S371F, S373P, S375F, T376A, D405N, R408S, K417N, S477N, T478K, E484A, Q493R, Q498R, N501Y, Y505H, D614G, H655Y, N679K, P681H, R682W, N764K, D796Y, Q954H, N969K), or sequence-verified Omicron BA.4 strain (alterations: V3G, T19I, Δ24-26, A27S, Δ69/70, G142D, V213G, G339D, S371F, S373P, S375F, T376A, D405N, R408S, K417N, N440K, L452R, S477N, T478K, E484A, F486V, Q498R, N501Y, Y505H, D614G, H655Y, N679K, P681H, N764K, D796Y, Q954H, N969K) to allow any antigen-specific antibodies to bind to the virus. A diagram of S glycoprotein alterations is shown in fig. S5b. The 2019-nCOV/ITALY-INMI1 strain S glycoprotein is identical in sequence to the wild-type SARS-CoV-2 S (Wuhan-Hu-1 isolate). Vero E6 (ATCC® CRL-1586™) cell monolayers were inoculated with the serum/virus mix in 96-well plates and incubated for 3 days (2019-nCOV/ITALY-INMI1 strain) or 4 days (Alpha, Beta, Delta, Omicron BA.1, BA.2 and BA.4 variant strain) to allow infection by non-neutralized virus. The plates were observed under an inverted light microscope and the wells were scored as positive for SARS-CoV-2 infection (i.e., showing CPE) or negative for SARS-CoV-2 infection (i.e., cells were alive without CPE). The neutralization titer was determined as the reciprocal of the highest serum dilution that protected more than 50% of cells from CPE and reported as GMT of duplicates. If no neutralization was observed, an arbitrary titer value of 5 (half of the LOD) was reported. Tables of the neutralization titers are provided (Tables S7 to S9).

### Depletion of RBD- or NTD-binding antibodies from human sera

SARS-CoV-2 wild-type strain S glycoprotein RBD- and NTD-coupled magnetic beads (Acro Biosystems, Cat.no. MBS-K002 and MBS-K019; 40 µg RBD/mg beads and 38 µg NTD/mg beads, respectively) were prepared according to the manufacturer’s instructions. Beads were resuspended in ultrapure water at 1 mg beads/mL and a magnet was used to collect and wash the beads with PBS. Beads were resuspended in serum to obtain 20 µg RBD- or NTD-bait per 100 µL serum. A mock depletion (undepleted control) was performed for each serum by adding 0.5 mg Biotin-saturated MyOne™ Streptavidin T1 Dynabeads™ (ThermoFisher, Cat.no. 65601) per 100 µL serum. Beads were incubated with human sera for 1 hour with gentle rotation. A magnet was used to separate bead-bound antibodies from the depleted supernatant. Depleted and undepleted sera were analyzed for cross-neutralization capacity using pseudovirus neutralization assays. Depletion efficacy for both RBD- and NTD-binding antibodies was determined by a multiplexed electrochemiluminescence immunoassay (Meso Scale Discovery, V-Plex SARS-CoV-2 Panel 1 Kit, Cat. No. K15359U-2). Table of the neutralization titers is provided (Table S11).

### Statistical analysis

The statistical method of aggregation used for the analysis of antibody titers is the geometric mean and for the ratio of SARS-CoV-2 VOC titer and wild-type strain titer the geometric mean and the corresponding 95% confidence interval. The use of the geometric mean accounts for the non-normal distribution of antibody titers, which span several orders of magnitude. The Friedman test with Dunn’s correction for multiple comparisons was used to conduct pairwise signed-rank tests of group geometric mean neutralizing antibody titers with a common control group. Spearman correlation was used to evaluate the monotonic relationship between non-normally distributed datasets. All statistical analyses were performed using GraphPad Prism software version 9.

## Acknowledgments

We thank the BioNTech global clinical Phase 2 trial (NCT04380701) participants, and the Omicron BA.1 and BA.2 convalescent research study participants from whom the post-immunization human sera were obtained. We thank the many colleagues at BioNTech and Pfizer who developed and produced the BNT162b2 vaccine candidate. We thank S. Jägle and N. Beckmann for logistical support; S. Shpyro, S. Nadim, C. Heiser, A. Telorman, C. Müller, A. Wanamaker, N. Williams, and J. VanCamp for sample demographics support; and the VisMederi team for work on live virus–neutralizing antibody assays.

## Funding

This work was supported by BioNTech.

## Author contributions

U.S., Ö.T., and A.M. conceived and conceptualized the work. A.M, and B.G.L. planned and supervised experiments. K.K., O.O., S.H., and S.C. coordinated and conducted sample collection. K.G. coordinated sample shipments and clinical data transfer. A.M., B.G.L., M.B., and A.W. performed experiments. A.M., and B.G.L. analyzed data. U.S., Ö.T., A.M., A.T., and A.F. interpreted data and wrote the manuscript. All authors supported the review of the manuscript.

## Competing interests

U.S. and Ö.T. are management board members and employees at BioNTech SE. A.M., B.G.L., K.K., A.W., M.B., A.F., A.T., and O.O. are employees at BioNTech SE. K.G., S.H. and S.C. are employees at University Hospital, Goethe University Frankfurt. U.G. is an employee at the Health Protection Authority, City of Frankfurt am Main. U.S., Ö.T. and A.M. are inventors on patents and patent applications related to RNA technology and COVID-19 vaccines. U.S., Ö.T., A.M., B.G.L., K.K., A.W., M.B., A.F., A.T., and O.O. have securities from BioNTech SE. S.C. has received honorarium for serving on a clinical advisory board for BioNTech.

## Data and materials availability

Participant baseline characteristics are provided in Table S1 to Table S3 and Table S10. The neutralization titers are provided in Tables S4 to S9 and Table S11.

Materials are available from the authors under a material transfer agreement with BioNTech.

---

**Fig. S1.**
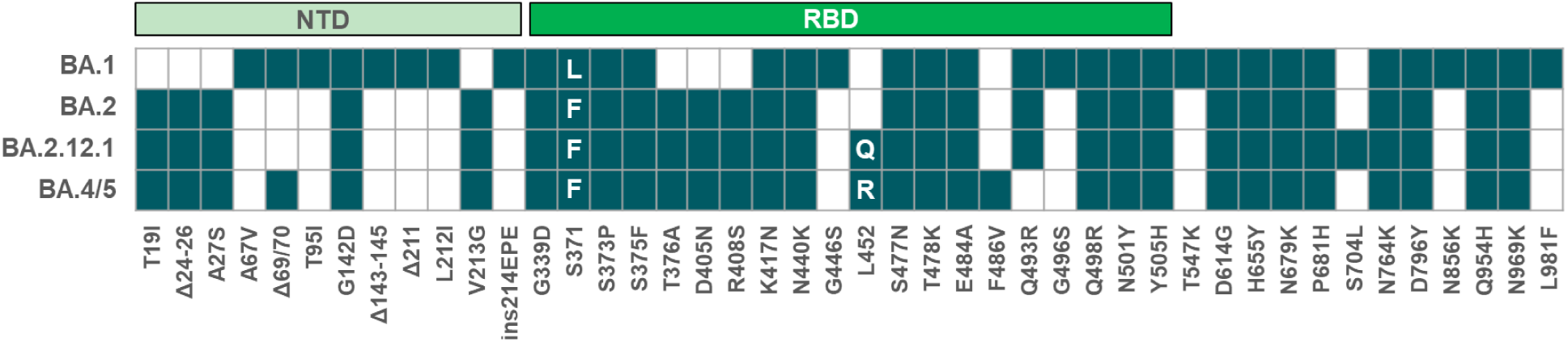
Alterations of the spike glycoprotein amino acid sequence of SARS-CoV-2 Omicron sub-lineages. Amino acid exchange and mutation type (substitutions, deletions, insertions) are indicated. White letters in boxes indicate the amino acid substitution per sub-lineage; Δ, deletion; ins, insertion; NTD, N-terminal domain; RBD, receptor-binding domain

**Fig. S2.**
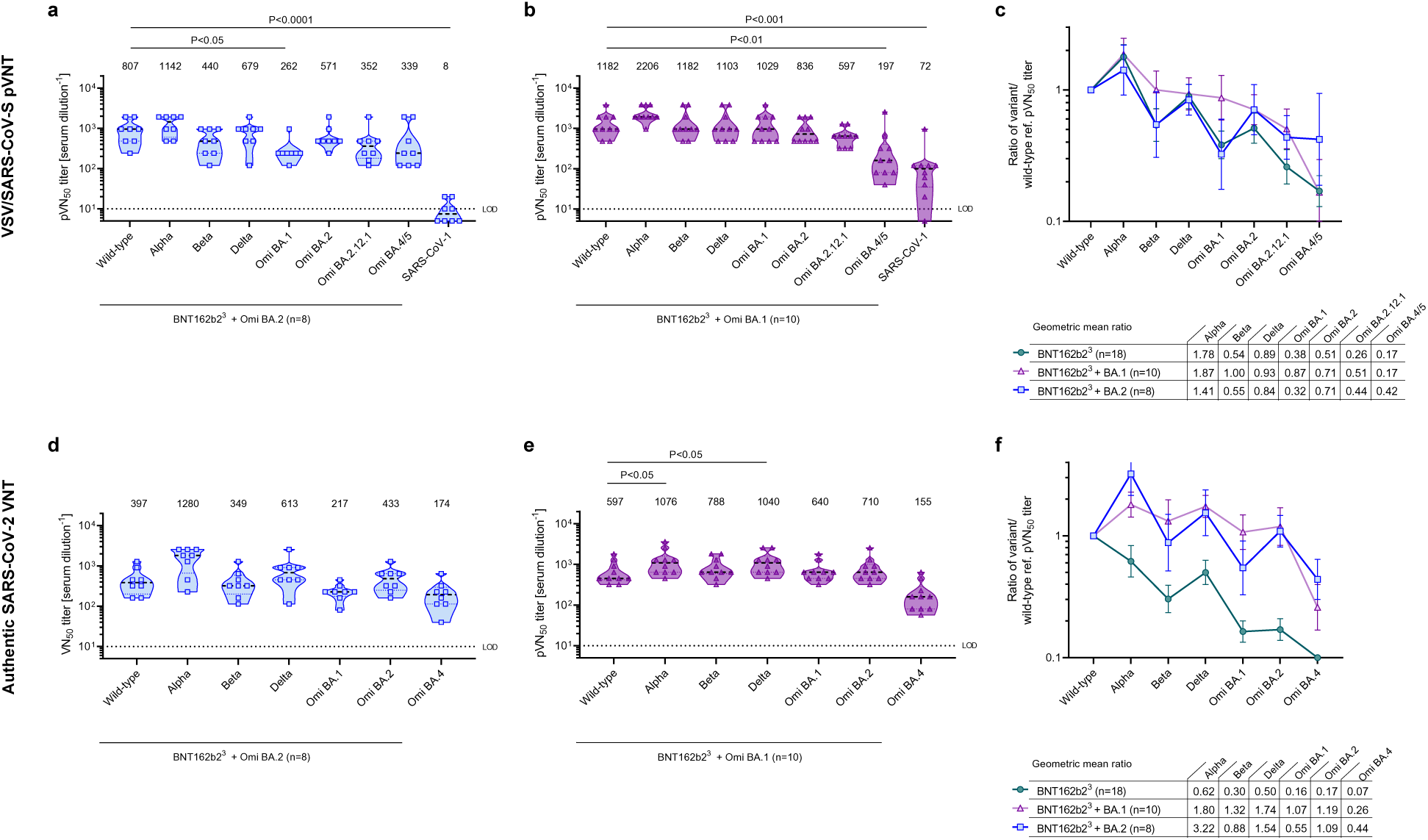
Omicron BA.2 breakthrough infection of BNT162b2 triple-vaccinated individuals induces broad neutralization of VOCs including Omicron BA.4/BA.5. Cohorts and serum sampling as described in Fig. 1. (a-b) 50% pseudovirus neutralization (pVN_50_) geometric mean titers (GMTs) against the indicated SARS-CoV-2 variants of concern (VOCs) or SARS-CoV-1 pseudoviruses. Values above violin plots represent group GMTs. (c) The ratio of SARS-CoV-2 VOC pVN_50_ GMTs normalized against the wild-type strain pVN_50_ GMT. Geometric mean ratios for the Omicron BA.2 breakthrough infected cohort were compared to data previously published in Quandt et al. (*1*) for BNT162b2^3^ and BNT162b2^3^ + BA.1, except for newly generated BA.2.12.1 neutralization data. Group geometric mean ratios with 95% confidence intervals are shown. (d-e) 50% virus neutralization (VN_50_) GMTs for BNT162b2^3^ + BA.1 and BNT162b2^3^ + BA.2. Values above violin plots represent group GMTs. (f) The ratio of SARS-CoV-2 VOC GMTs normalized against the wild-type strain VN_50_ GMT. Serum was tested in duplicate. For titer values below the limit of detection (LOD), LOD/2 values are plotted. The non-parametric Friedman test with Dunn’s multiple comparisons correction was used to compare the group GMT against the wild-type strain with group GMTs against the indicated variants and SARS-CoV-1. Multiplicity-adjusted p values are shown.

**Fig. S3.**
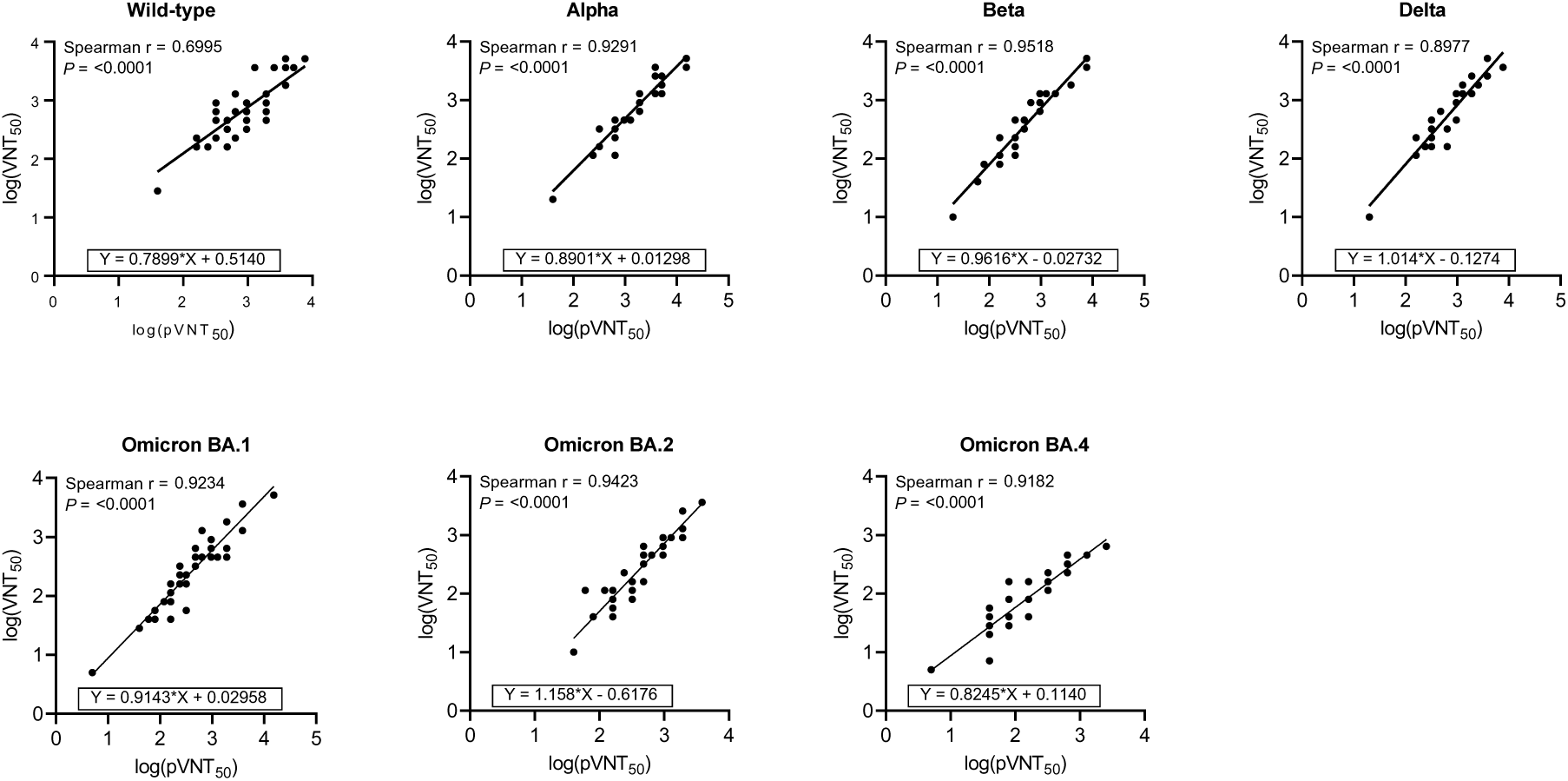
50% pseudovirus neutralization (pVN_50_) correlates with 50% live SARS-CoV-2 neutralization (VN_50_) titer data. Nonparametric Spearman correlation of VSV-SARS-CoV-2 pVN_50_ with live SARS-CoV-2 VN_50_ titers for n=45 serum samples drawn from SARS-CoV-2-naïve BNT162b2 triple-vaccinated individuals (BNT162b2^3^; n=18) after the third dose, from triple mRNA vaccinated individuals with subsequent Omicron BA.1 breakthrough infection (mRNA-Vax^3^ + BA.1; n=14) post-infection, and from triple mRNA vaccinated individuals with subsequent Omicron BA.2 breakthrough infection (mRNA-Vax^3^ + BA.2; n=13) post-infection. Correlations are plotted per SARS-CoV-2 variant. Correlation coefficient r, two-tailed P values and the linear equation are given.

**Fig. S4.**
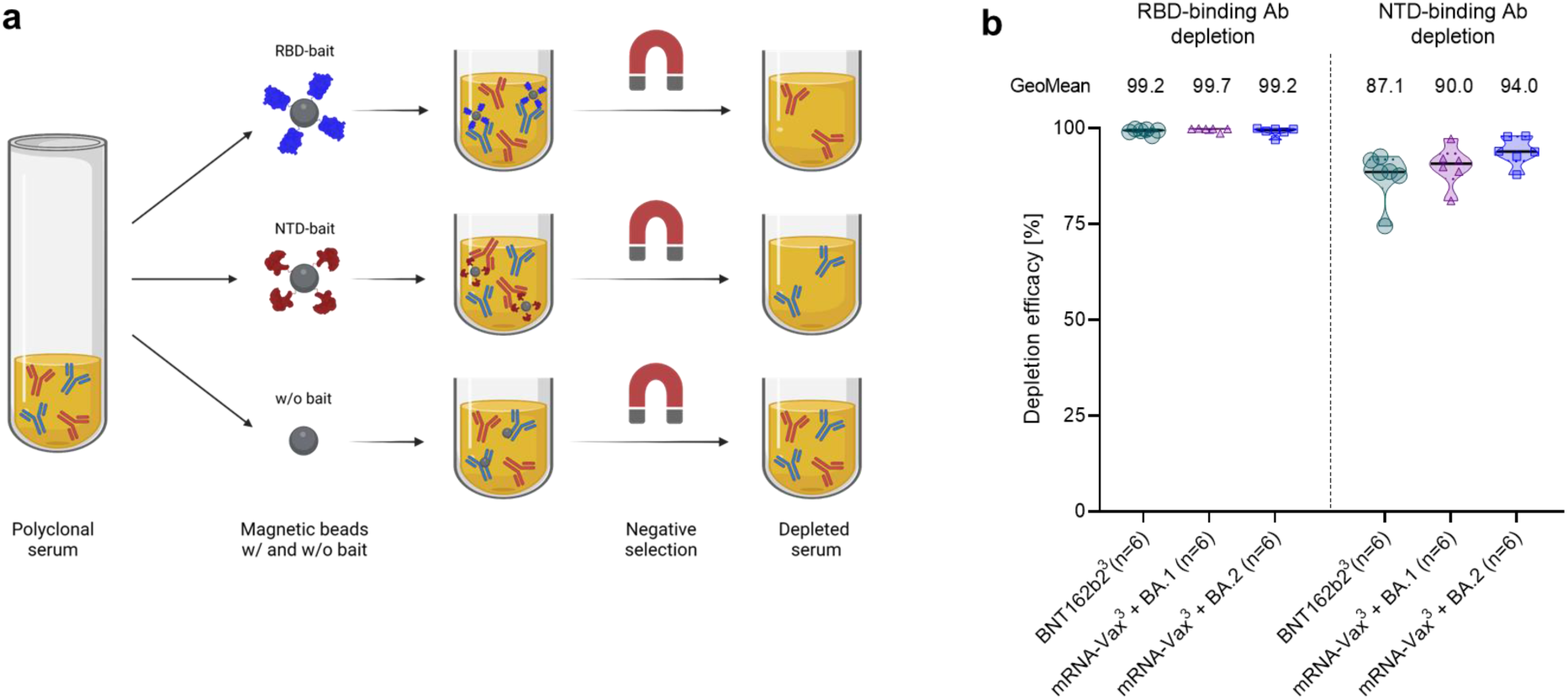
RBD-binding and NTD-binding antibodies can be depleted from human serum. Serum was drawn from SARS-CoV-2-naïve BNT162b2 triple-vaccinated individuals (BNT162b2^3^; n=6), and from triple mRNA vaccinated individuals with Omicron BA.1 (mRNA-Vax^3^ + BA.1; n=6) or Omicron BA.2 breakthrough infection (mRNA-Vax^3^ + BA.2; n=6). Magnetic bead technology was used for depleting serum of RBD- or NTD-binding antibodies, or for mock depleting. (a) Schematic of antibody depletion from serum. (b) The relative concentration of RBD-binding and NTD-binding antibodies was determined by a multiplexed electrochemiluminescence immunoassay. The relative decrease in antibody concentrations in depleted compared to mock-depleted sera are shown. Numbers above graph depict geometric mean reduction within groups.

**Fig. S5.**
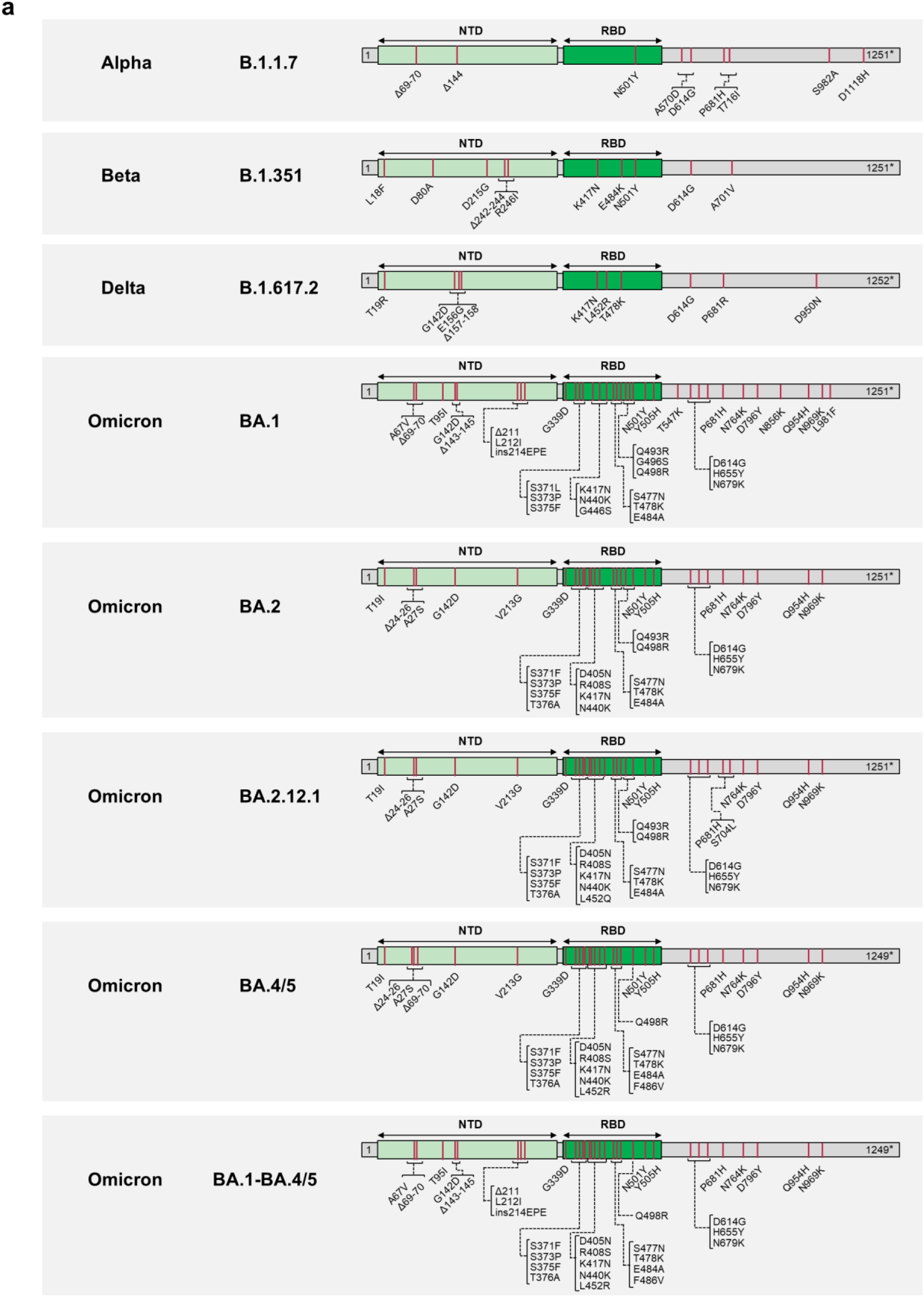
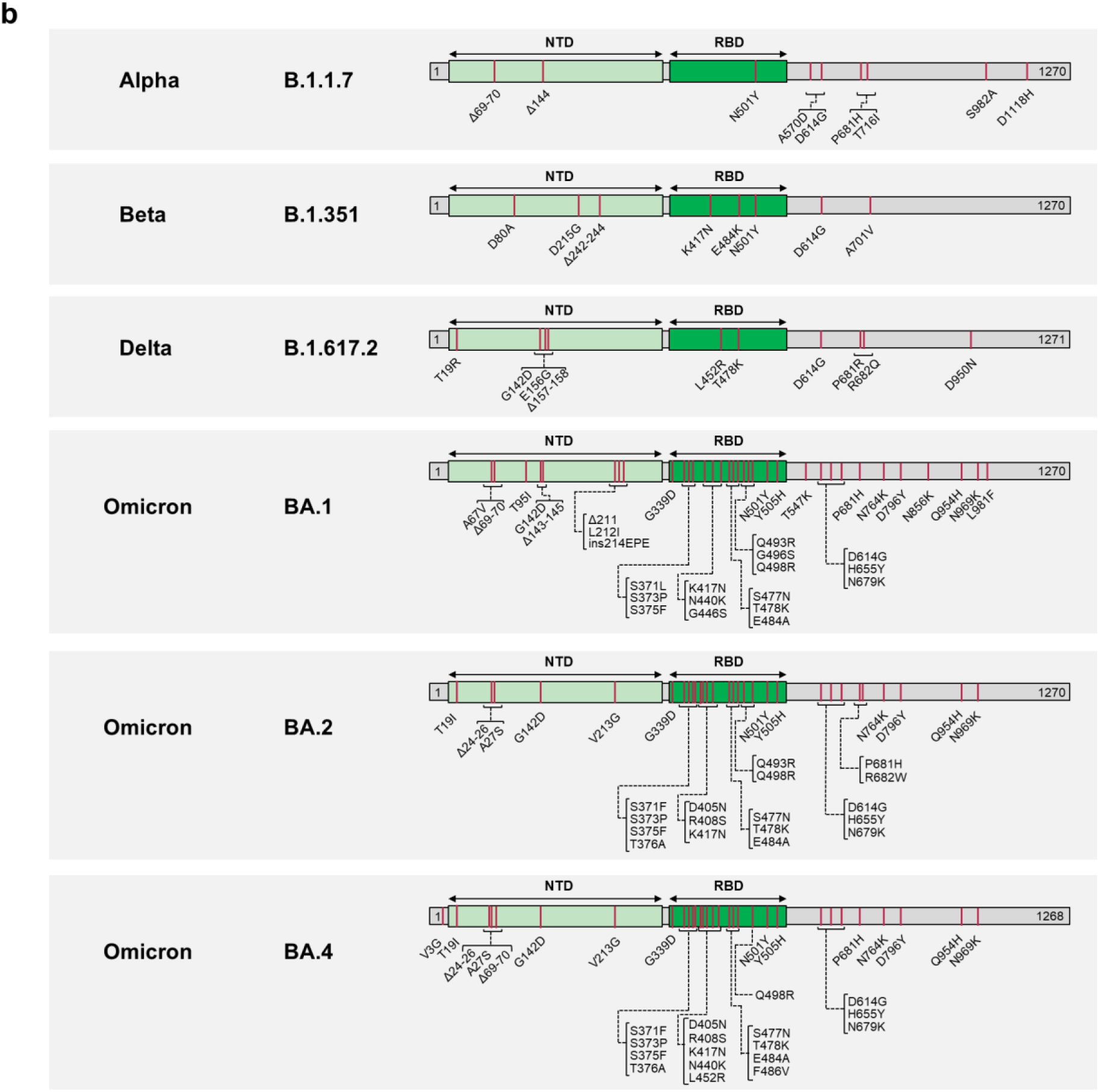
Characterization of SARS-CoV-2 S glycoproteins used in the assays based on (a) VSV-SARS-CoV-2 variant pseudoviruses and (b) live authentic SARS-CoV-2. The sequence of the Wuhan-Hu-1 isolate SARS-CoV-2 S glycoprotein (GenBank: QHD43416.1) was used as reference. Amino acid positions, amino acid descriptions (one letter code) and kind of alterations (substitutions, deletions, insertions) are indicated. NTD, N-terminal domain; RBD, Receptor-binding domain, Δ, deletion; ins, insertion; *, Cytoplasmic domain truncated for the C-terminal 19 amino acids.

**Table S1.** Vaccinated individuals analyzed for neutralizing antibody responses.

| Characteristic | BNT162b2<sup>3</sup><br>(n=18) | mRNA-Vax<sup>3</sup><br>+ BA.1<br>(n=14) | mRNA-Vax<sup>3</sup><br>+ BA.2<br>(n=13) |
| --- | --- | --- | --- |
| Sex, n (%) |  |  |  |
| Male | 9 (50) | 11 (79) | 4 (31) |
| Female | 9 (50) | 3 (21) | 9 (69) |
| Age, median (range) | 38 (23-54) | 32 (23-60) | 30 (25-72) |
| Age group at vaccination, n (%) |  |  |  |
| 18-55 yrs | 18 (100) | 12 (86) | 11 (85) |
| 56-85 yrs | 0 (0) | 2 (14) | 2 (15) |
| SARS-CoV-2 status, n (%) |  |  |  |
| Positive | 0 (0) | 14 (100)<sup>#</sup> | 13 (100)* |
| Negative | 18 (100)<sup>†</sup> | 0 (0) | 0 (0) |
| Unknown | 0 (0) | 0 (0) | 0 (0) |
| Interval, median (range) |  |  |  |
| Days between D1/D2 | ‡ | 38 (20-92) | 42 (15-43) |
| Days between D2/D3 | 202 (181-266) | 192 (154-256) | 184 (152-259) |
| Days until serum draw after D3 | 28 (26-30) | N/A | N/A |
| Days between last dose/infection | N/A | 25 (3-112) | 135 (36-200) |
| Days until serum draw after infection | N/A | 43 (25-55) | 39 (30-99) |

N/A, not applicable; D, dose; yrs, years; n, number.
\*, Individuals experienced SARS-CoV-2 breakthrough infections between March and May 2022, during which period the BA.2 lineage was dominant in Germany
<sup>#</sup>, Omicron infection PCR-confirmed at time of recruitment to the research study. Individuals experienced SARS-CoV-2 breakthrough infections between November 2021 and January 2022, during which period the BA.1 lineage was dominant in Germany
<sup>†</sup>, No evidence of prior SARS-CoV-2 infection (based on COVID-19 symptoms/signs and SARS-CoV-2 PCR test)
<sup>‡</sup>, Participants received the primary 2-dose series of BNT162b2 vaccine as part of a governmental vaccination program and the interval between doses was not recorded

**Table S2.** Individuals triple vaccinated with mRNA COVID-19 vaccine and subsequently infected with Omicron BA.1 (mRNA-Vax^3^ + BA.1).

| Participant ID | Age | Sex | Vaccination | Date positive test | Omicron subtype | Dose 1-2 interval (days) | Dose 2-3 interval (days) | Positive test after last vaccination (days) | Blood draw after positive test (days) | Severity (WHO grade) |
| --- | --- | --- | --- | --- | --- | --- | --- | --- | --- | --- |
| 14 | 32 | f | BNT<sup>3</sup> | NOV2021 | BA.1 | 24 | 243 | 64 | 55 | 1-2 |
| 15 | 32 | m | BNT<sup>3</sup> | NOV2021 | BA.1 | 20 | 233 | 66 | 53 | 1-2 |
| 16 | 28 | m | BNT<sup>3</sup> | DEC2021 | n/a | 35 | 213 | 10 | 47 | 1-2 |
| 17 | 29 | f | BNT<sup>3</sup> | DEC2021 | BA.1 | 36 | 189 | 3 | 46 | 1-2 |
| 18 | 23 | m | BNT<sup>3</sup> | DEC2021 | n/a | 42 | 159 | 27 | 44 | 1-2 |
| 19 | 31 | m | BNT<sup>3</sup> | DEC2021 | n/a | 42 | 166 | 20 | 43 | 1-2 |
| 20 | 53 | m | BNT<sup>3</sup> | JAN2022 | n/a | 39 | 194 | 22 | 25 | 1-2 |
| 21 | 50 | f | BNT<sup>3</sup> | JAN2022 | n/a | 92 | 169 | 35 | 28 | 1-2 |
| 22 | 50 | m | BNT<sup>3</sup> | JAN2022 | n/a | 42 | 169 | 44 | 31 | 1-2 |
| 23 | 60 | m | BNT<sup>3</sup> | JAN2022 | n/a | 26 | 236 | 112 | 43 | 1-2 |
| 24 | 28 | m | MOD<sup>2</sup>/BNT | DEC2021 | n/a | 28 | 252 | 22 | 40 | 1-2 |
| 25 | 32 | m | MOD<sup>2</sup>/BNT | DEC2021 | n/a | 42 | 154 | 13 | 42 | 1-2 |
| 26 | 50 | m | MOD<sup>2</sup>/BNT | JAN2022 | n/a | 28 | 256 | 45 | 31 | 1-2 |
| 27 | 60 | m | MOD<sup>3</sup> | DEC2021 | BA.1 | 42 | 172 | 3 | 35 | 1-2 |
| **Median** | **32** | **N/A** | **N/A** | **N/A** | **N/A** | **38** | **192** | **25** | **43** | **N/A** |

m, male; f, female; n/a, not available; N/A, not applicable
BNT, BioNTech/Pfizer BNT162b2; MOD, Moderna mRNA-1273; BNT<sup>3</sup>, BNT162b2 three-dose series; MOD<sup>2</sup>, mRNA-1273 two-dose series; MOD<sup>3</sup>, mRNA-1273 three-dose series

**Table S3.** Individuals triple vaccinated with mRNA COVID-19 vaccine and subsequently infected with Omicron BA.2 (mRNA-Vax^3^ + BA.2).

| Participant ID | Age | Sex | Vaccination | Date positive test | Omicron subtype | Dose 1-2 interval (days) | Dose 2-3 interval (days) | Positive test after last vaccination (days) | Blood draw after positive test (days) | Severity (WHO grade) |
| --- | --- | --- | --- | --- | --- | --- | --- | --- | --- | --- |
| 1 | 42 | F | BNT<sup>3</sup> | APR2022 | n/a | 42 | 158 | 135 | 36 | 1-2 |
| 2 | 44 | M | BNT<sup>2</sup>/MOD | APR2022 | n/a | 42 | 164 | 129 | 36 | 1-2 |
| 3 | 28 | f | MOD<sup>2</sup>/BNT | MAR2022 | n/a | 42 | 163 | 36 | 99 | 1-2 |
| 4 | 26 | f | BNT<sup>3</sup> | APR2022 | n/a | 43 | 152 | 104 | 58 | 1-2 |
| 5 | 34 | m | BNT<sup>3</sup> | MAY2022 | n/a | 40 | 154 | 141 | 37 | 1-2 |
| 6 | 29 | m | BNT<sup>3</sup> | APR2022 | n/a | 42 | 165 | 89 | 63 | 1-2 |
| 7 | 57 | f | MOD<sup>2</sup>/BNT | APR2022 | n/a | 15 | 244 | 153 | 46 | 1-2 |
| 8 | 25 | m | MOD/BNT<sup>2</sup> | MAY2022 | n/a | 28 | 217 | 188 | 34 | 1-2 |
| 9 | 30 | f | MOD<sup>2</sup>/BNT | APR2022 | n/a | 42 | 195 | 129 | 66 | 1-2 |
| 10 | 28 | f | BNT<sup>2</sup>/MOD | MAY2022 | n/a | 21 | 224 | 64 | 39 | 1-2 |
| 11 | 29 | f | BNT<sup>3</sup> | APR2022 | n/a | 22 | 249 | 149 | 64 | 1-2 |
| 12 | 53 | f | BNT<sup>3</sup> | MAY2022 | n/a | 29 | 259 | 200 | 30 | 1-2 |
| 13 | 72 | f | BNT<sup>3</sup> | MAY2022 | n/a | 42 | 184 | 157 | 30 | 1-2 |
| **Median** | **30** | **N/A** | **N/A** | **N/A** | **N/A** | **42** | **184** | **135** | **39** | **N/A** |

m, male; f, female; n/a, not available; N/A, not applicable
BNT, BioNTech/Pfizer BNT162b2; MOD, Moderna mRNA-1273; BNT<sup>2</sup>, BNT162b2 two-dose series; BNT<sup>3</sup>, BNT162b2 three-dose series; MOD<sup>2</sup>, mRNA-1273 two-dose series

**Table S4.** pVN_50_ values of sera collected from SARS-CoV-2-naïve triple-vaccinated individuals (BNT162b2^3^)

| Participant ID | pVN<sub>50</sub> |  |  |  |  |  |  |  |  | SARS-CoV-1 |
| --- | --- | --- | --- | --- | --- | --- | --- | --- | --- | --- |
|  | Wild-type | Alpha | Beta | Delta | Omicron BA.1 | Omicron BA.2 | Omicron BA.2.12.1 | Omicron BA.4/5 | Omicron BA.1-BA.4/5 |  |
| 28 | 160 | 320 | 160 | 160 | 80 | 160 | 80 | 40 | 40 | 5 |
| 29 | 640 | 640 | 320 | 320 | 160 | 320 | 160 | 40 | 40 | 40 |
| 30 | 5120 | 5120 | 1280 | 2560 | 1280 | 1280 | 640 | 640 | 160 | 40 |
| 31 | 320 | 640 | 160 | 320 | 160 | 160 | 80 | 40 | 40 | 20 |
| 32 | 640 | 640 | 80 | 640 | 320 | 160 | 40 | 40 | 20 | 20 |
| 33 | 320 | 640 | 160 | 320 | 160 | 160 | 80 | 40 | 20 | 10 |
| 34 | 320 | 640 | 320 | 320 | 160 | 160 | 80 | 80 | 40 | 10 |
| 35 | 320 | 640 | 320 | 320 | 160 | 160 | 160 | 80 | 40 | 20 |
| 36 | 160 | 320 | 80 | 160 | 40 | 80 | 40 | 40 | 10 | 20 |
| 37 | 320 | 1280 | 160 | 320 | 160 | 60 | 160 | 80 | 40 | 20 |
| 38 | 1280 | 5120 | 640 | 1280 | 640 | 640 | 640 | 320 | 160 | 80 |
| 39 | 40 | 40 | 20 | 20 | 5 | 40 | 10 | 5 | 5 | 5 |
| 40 | 320 | 640 | 320 | 320 | 80 | 160 | 80 | 40 | 20 | 20 |
| 41 | 160 | 320 | 160 | 320 | 80 | 160 | 80 | 40 | 20 | 20 |
| 42 | 320 | 640 | 320 | 320 | 320 | 320 | 160 | 160 | 40 | 20 |
| 43 | 640 | 640 | 320 | 320 | 160 | 320 | 80 | 80 | 40 | 40 |
| 44 | 2560 | 5120 | 640 | 1280 | 640 | 640 | 320 | 320 | 160 | 80 |
| 45 | 320 | 640 | 160 | 640 | 160 | 320 | 80 | 80 | 80 | 20 |

**Table S5.** pVN_50_ values of sera collected from individuals with Omicron BA.1 breakthrough infection (mRNA-Vax^3^ + BA.1)

| Participant ID | pVN<sub>50</sub> |  |  |  |  |  |  |  |  | SARS-CoV-1 |
| --- | --- | --- | --- | --- | --- | --- | --- | --- | --- | --- |
|  | Wild-type | Alpha | Beta | Delta | Omicron BA.1 | Omicron BA.2 | Omicron BA.2.12.1 | Omicron BA.4/5 | Omicron BA.1-BA.4/5 |  |
| 14 | 1920 | 3840 | 1920 | 1920 | 1920 | 960 | 640 | 320 | 80 | 120 |
| 15 | 960 | 1920 | 960 | 480 | 480 | 480 | 640 | 160 | 320 | 120 |
| 16 | 3840 | 3840 | 3840 | 3840 | 1920 | 1920 | 1280 | 640 | 640 | 960 |
| 17 | 960 | 1920 | 960 | 960 | 960 | 960 | 640 | 160 | 160 | 120 |
| 18 | 480 | 1920 | 960 | 480 | 480 | 480 | 640 | 40 | 40 | 20 |
| 19 | 1920 | 1920 | 960 | 960 | 960 | 1920 | 640 | 320 | 320 | 40 |
| 20 | 960 | 1920 | 960 | 960 | 1920 | 480 | 320 | 80 | 160 | 80 |
| 21 | 960 | 1920 | 480 | 960 | 480 | 480 | 320 | 80 | 80 | 5 |
| 22 | 480 | 960 | 480 | 480 | 480 | 480 | 320 | 80 | 40 | 60 |
| 23 | 1920 | 3840 | 3840 | 3840 | 3840 | 1920 | 1280 | 2560 | 1280 | 120 |
| 24 | 3840 | 15360 | 7680 | 7680 | 3840 | 3840 | 2560 | 2560 | 1280 | 120 |
| 25 | 7680 | 15360 | 7680 | 3840 | 15360 | 3840 | 2560 | 1280 | 640 | 480 |
| 26 | 480 | 240 | 60 | 240 | 60 | 120 | 40 | 40 | 10 | 30 |
| 27 | 1920 | 1920 | 960 | 1920 | 960 | 960 | 640 | 640 | 320 | 60 |

**Table S6.**
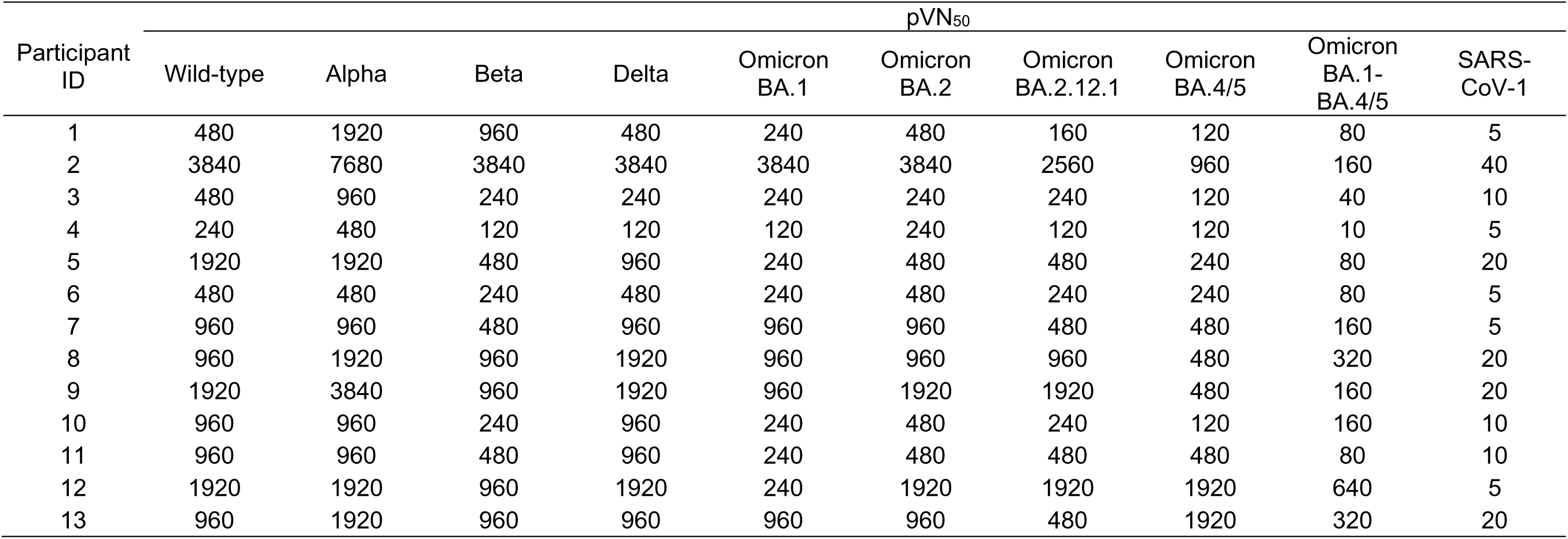
pVN_50_ values of sera collected from individuals with Omicron BA.2 breakthrough infection (mRNA-Vax^3^ + BA.2)

**Table S7.** VN_50_ values of sera collected from SARS-CoV-2-naïve triple-vaccinated individuals (BNT162b2^3^)

| Participant ID | VN<sub>50</sub> |  |  |  |  |  |  |
| --- | --- | --- | --- | --- | --- | --- | --- |
|  | Wild-type | Alpha | Beta | Delta | Omicron BA.1 | Omicron BA.2 | Omicron BA.4 |
| 28 | 226 | 320 | 80 | 226 | 40 | 40 | 28 |
| 29 | 640 | 320 | 226 | 226 | 160 | 80 | 28 |
| 30 | 3620 | 2560 | 1280 | 1810 | 453 | 905 | 226 |
| 31 | 226 | 113 | 226 | 160 | 40 | 80 | 28 |
| 32 | 226 | 453 | 80 | 160 | 57 | 40 | 20 |
| 33 | 640 | 453 | 113 | 226 | 80 | 57 | 40 |
| 34 | 453 | 226 | 226 | 453 | 80 | 80 | 28 |
| 35 | 640 | 320 | 226 | 226 | 160 | 113 | 40 |
| 36 | 160 | 160 | 80 | 113 | 28 | 40 | 20 |
| 37 | 640 | 453 | 80 | 320 | 80 | 113 | 40 |
| 38 | 3620 | 1280 | 905 | 1280 | 453 | 453 | 160 |
| 38 | 28 | 20 | 10 | 10 | 5 | 10 | 5 |
| 39 | 453 | 320 | 113 | 226 | 40 | 80 | 20 |
| 40 | 226 | 320 | 113 | 226 | 57 | 57 | 28 |
| 41 | 905 | 226 | 160 | 453 | 160 | 113 | 40 |
| 42 | 1280 | 226 | 160 | 226 | 113 | 113 | 40 |
| 43 | 3620 | 1810 | 905 | 1810 | 1280 | 453 | 160 |
| 44 | 905 | 453 | 226 | 320 | 80 | 113 | 40 |
| 45 | 226 | 320 | 80 | 226 | 40 | 40 | 28 |

**Table S8.**
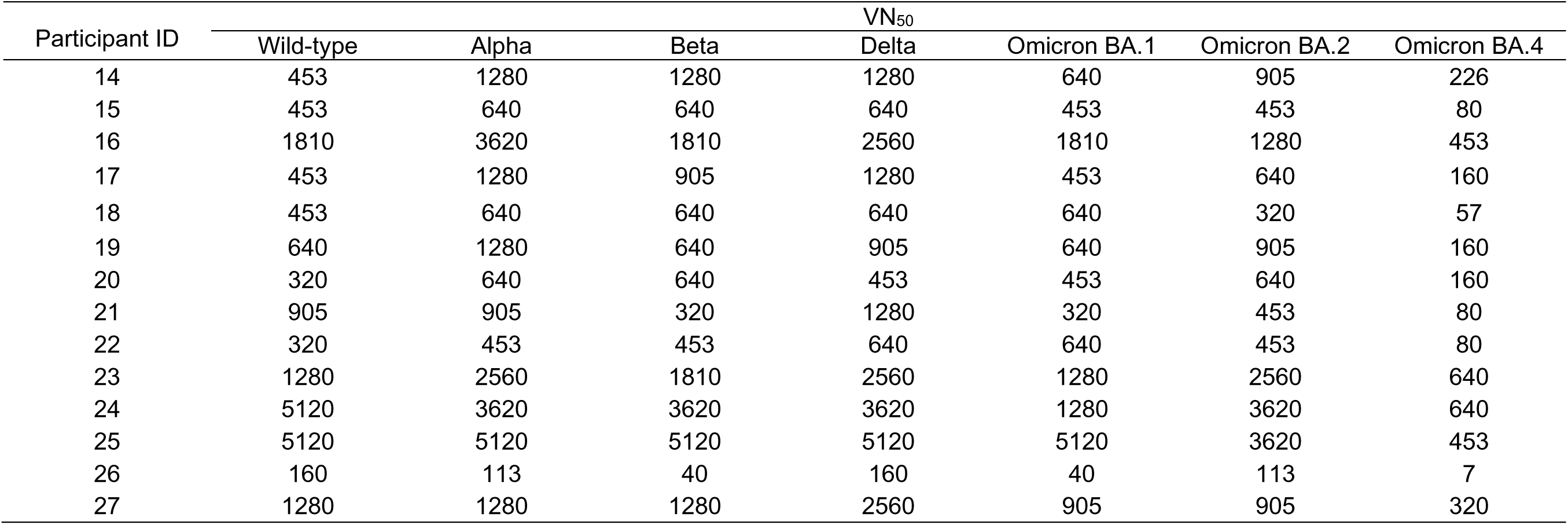
VN_50_ values of sera collected from individuals with Omicron BA.1 breakthrough infection (mRNA-Vax^3^ + BA.1)

**Table S9.** VN_50_ values of sera collected from individuals with Omicron BA.2 breakthrough infection (mRNA-Vax^3^ + BA.2)

| Participant ID | VN<sub>50</sub> |  |  |  |  |  |  |
| --- | --- | --- | --- | --- | --- | --- | --- |
|  | Wild-type | Alpha | Beta | Delta | Omicron BA.1 | Omicron BA.2 | Omicron BA.4 |
| 1 | 320 | 1810 | 320 | 453 | 226 | 320 | 113 |
| 2 | 3620 | 5120 | 3620 | 5120 | 3620 | 3620 | 905 |
| 3 | 320 | 905 | 226 | 226 | 160 | 226 | 113 |
| 4 | 160 | 226 | 113 | 113 | 80 | 226 | 40 |
| 5 | 905 | 2560 | 320 | 905 | 226 | 640 | 226 |
| 6 | 160 | 453 | 160 | 453 | 226 | 160 | 113 |
| 7 | 640 | 1810 | 320 | 640 | 640 | 640 | 226 |
| 8 | 640 | 1810 | 1810 | 1280 | 905 | 453 | 226 |
| 9 | 905 | 1810 | 2560 | 2560 | 640 | 1280 | 320 |
| 10 | 453 | 1810 | 320 | 453 | 226 | 320 | 160 |
| 11 | 320 | 1280 | 453 | 905 | 160 | 640 | 226 |
| 12 | 1280 | 2560 | 640 | 2560 | 320 | 1280 | 640 |
| 13 | 453 | 2560 | 1280 | 905 | 453 | 640 | 320 |

**Table S10.** Vaccinated individuals analyzed for neutralizing antibody responses after depletion of RBD-/NTD-binding antibodies.

| Characteristic | mRNA-Vax<sup>3</sup><br>+ BA.2<br>(n=6) | mRNA-Vax<sup>3</sup><br>+ BA.1<br>(n=6) | BNT162b2<sup>3</sup><br>(n=6) |
| --- | --- | --- | --- |
| Sex, n (%) |  |  |  |
| Male | 1 (17) | 4 (67) | 4 (67) |
| Female | 5 (83) | 2 (33) | 2 (33) |
| Age, median (range) | 41.5 (25-72) | 32 (29-53) | 36 (23-49) |
| Age group at vaccination, n (%) |  |  |  |
| 18-55 yrs | 4 (67) | 6 (100) | 6 (100) |
| 56-85 yrs | 2 (33) | 0 (0) | 0 (0) |
| SARS-CoV-2 status, n (%) |  |  |  |
| Positive | 6 (100)* | 6 (100)<sup>#</sup> | 0 (0) |
| Negative | 0 (0) | 0 (0) | 6 (100)<sup>†</sup> |
| Unknown | 0 (0) | 0 (0) | 0 (0) |
| Interval, median (range) |  |  |  |
| Days between D1/D2 | 28.5 (15-42) | 39 (20-92) | ‡ |
| Days between D2/D3 | 230.5 (184-259) | 189 (154-233) | 219 (181-266) |
| Days until serum draw after D3 | N/A | N/A | 27.5 (26-30) |
| Days between last dose/infection | 155 (129-200) | 20 (3-66) | N/A |
| Days until serum draw after infection | 40 (30-66) | 43 (25-53) | N/A |

N/A: not applicable; D, Dose; Yrs, Years; n, Number.
\*, Individuals experienced SARS-CoV-2 breakthrough infections between March and May 2022, during which period the BA.2 lineage was dominant in Germany
<sup>#</sup>, Omicron infection PCR-confirmed at time of recruitment to the research study. Individuals experienced SARS-CoV-2 breakthrough infections between November 2021 and January 2022, during which period the BA.1 lineage was dominant in Germany
<sup>†</sup>, No evidence of prior SARS-CoV-2 infection (based on COVID-19 symptoms/signs and SARS-CoV-2 PCR test)
<sup>‡</sup>, Participants received the primary 2-dose series of BNT162b2 vaccine as part of a governmental vaccination program and the interval between doses was not recorded.

**Table S11.** pVN_50_ values of sera depleted of NTD or RBD-binding antibodies

| Participant ID | Cohort | pVN<sub>50</sub> |  |  |  |  |  |  |  |  |  |  |  |
| --- | --- | --- | --- | --- | --- | --- | --- | --- | --- | --- | --- | --- | --- |
|  |  | Wild-type |  |  | Omicron BA.1 |  |  | Omicron BA.2 |  |  | Omicron BA.4/5 |  |  |
|  |  | Mock | NTD | RBD | Mock | NTD | RBD | Mock | NTD | RBD | Mock | NTD | RBD |
| 7 | mRNA-Vax<sup>3</sup>+BA.2 | 1179 | 1021 | 200 | 996 | 1143 | 10 | 880 | 668 | 219 | 494 | 252 | 193 |
| 8 | mRNA-Vax<sup>3</sup>+BA.2 | 2039 | 1187 | 158 | 1276 | 1491 | 10 | 981 | 730 | 162 | 1148 | 525 | 123 |
| 9 | mRNA-Vax<sup>3</sup>+BA.2 | 2226 | 1607 | 673 | 1224 | 1276 | 10 | 1217 | 738 | 460 | 1139 | 420 | 535 |
| 11 | mRNA-Vax<sup>3</sup>+BA.2 | 1435 | 939 | 251 | 650 | 641 | 10 | 537 | 346 | 169 | 538 | 273 | 97 |
| 12 | mRNA-Vax<sup>3</sup>+BA.2 | 3562 | 2287 | 834 | 674 | 479 | 10 | 5562 | 805 | 1129 | 1797 | 626 | 1043 |
| 13 | mRNA-Vax<sup>3</sup>+BA.2 | 1463 | 2956 | 98 | 1401 | 1219 | 10 | 1403 | 732 | 247 | 821 | 634 | 118 |
| 15 | mRNA-Vax<sup>3</sup>+BA.1 | 1037 | 953 | 103 | 601 | 623 | 10 | 359 | 354 | 10 | 239 | 232 | 10 |
| 17 | mRNA-Vax<sup>3</sup>+BA.1 | 1401 | 1681 | 61 | 1525 | 1379 | 10 | 973 | 740 | 25 | 492 | 430 | 45 |
| 19 | mRNA-Vax<sup>3</sup>+BA.1 | 1992 | 1589 | 235 | 1754 | 1363 | 54 | 990 | 923 | 10 | 573 | 438 | 10 |
| 20 | mRNA-Vax<sup>3</sup>+BA.1 | 842 | 766 | 127 | 789 | 836 | 10 | 825 | 557 | 10 | 241 | 319 | 10 |
| 21 | mRNA-Vax<sup>3</sup>+BA.1 | 1404 | 1751 | 77 | 697 | 546 | 10 | 388 | 556 | 10 | 277 | 254 | 10 |
| 25 | mRNA-Vax<sup>3</sup>+BA.1 | 1870 | 1224 | 219 | 1439 | 1613 | 46 | 849 | 1216 | 27 | 886 | 612 | 10 |
| 30 | BNT162b2<sup>3</sup> | 3818 | 2279 | 497 | 746 | 923 | 10 | 1349 | 644 | 463 | 620 | 238 | 278 |
| 32 | BNT162b2<sup>3</sup> | 521 | 537 | 160 | 143 | 173 | 10 | 98 | 644 | 10 | 59 | 238 | 10 |
| 38 | BNT162b2<sup>3</sup> | 2085 | 2332 | 348 | 1014 | 1147 | 39 | 844 | 931 | 38 | 364 | 276 | 56 |
| 43 | BNT162b2<sup>3</sup> | 631 | 671 | 281 | 237 | 265 | 10 | 187 | 164 | 40 | 108 | 101 | 36 |
| 44 | BNT162b2<sup>3</sup> | 1985 | 2071 | 463 | 745 | 591 | 10 | 525 | 648 | 61 | 247 | 222 | 47 |
| 45 | BNT162b2<sup>3</sup> | 631 | 487 | 133 | 191 | 167 | 10 | 131 | 130 | 10 | 119 | 97 | 10 |

## Notes

### Competing Interest Statement

U.S. and O.T. are management board members and employees at BioNTech SE. A.M., B.G.L., K.K., A.W., M.B., A.F., A.T., and O.O. are employees at BioNTech SE. K.G., S.H. and S.C. are employees at University Hospital, Goethe University Frankfurt. U.G. is an employee at the Health Protection Authority, City of Frankfurt am Main. U.S., O.T. and A.M. are inventors on patents and patent applications related to RNA technology and COVID-19 vaccines. U.S., O.T., A.M., B.G.L., K.K., A.W., M.B., A.F., A.T., and O.O. have securities from BioNTech SE. S.C. has received honorarium for serving on a clinical advisory board for BioNTech.

